# Multi-omics profiling establishes the polypharmacology of FDA Approved CDK4/6 inhibitors and the potential for differential clinical activity

**DOI:** 10.1101/211680

**Authors:** Marc Hafner, Caitlin E. Mills, Kartik Subramanian, Chen Chen, Mirra Chung, Sarah A. Boswell, Robert A. Everley, Changchang Liu, Charlotte S. Walmsley, Dejan Juric, Peter K. Sorger

## Abstract

The target profiles of many drugs are established early in their development and are not systematically revisited at the time of FDA approval. Thus, it is often unclear whether therapeutics with the same nominal targets but different chemical structures are functionally equivalent. In this paper we use five different phenotypic and biochemical assays to compare approved inhibitors of cyclin-dependent kinases 4/6 – collectively regarded as breakthroughs in the treatment of hormone receptor-positive breast cancer. We find that transcriptional, proteomic and phenotypic changes induced by palbociclib, ribociclib, and abemaciclib differ significantly; abemaciclib in particular has advantageous activities partially overlapping those of alvocidib, an older polyselective CDK inhibitor. In cells and mice, abemaciclib inhibits kinases other than CDK4/6 including CDK2/Cyclin A/E – implicated in resistance to CDK4/6 inhibition – and CDK1/Cyclin B. The multi-faceted experimental and computational approaches described here therefore uncover under-appreciated differences in CDK4/6 inhibitor activities with potential importance in treating human patients.

## INTRODUCTION

Progression through the cell cycle is controlled by more than a dozen distinct protein complexes involving cyclins and cyclin-dependent kinases (CDKs). Because dysregulation of the cell cycle is a hallmark of cancer, several generations of CDK inhibitors have been tested as potential therapeutic agents. However, identifying CDK inhibitors that are more active on tumor than normal cells has been a challenge and it is only recently that CDK4/6 inhibitors have emerged as promising therapies, particularly in breast cancer. CDK4 and CDK6 bind cyclin D early in the G1 phase of the cell cycle and phosphorylate the retinoblastoma protein (pRb). pRb is then hyper-phosphorylated by CDK2/cyclin E, relieving its inhibitory activities against transcription factors of the E2F family and allowing for S phase entry. Later in the cell cycle, CDK2/cyclin A and CDK1 in complex with cyclin A and B promote entry and progression through G2 and mitosis. Multiple genetic changes in cancer cells disrupt critical steps in cell cycle regulation: amplification of CDK4, CDK6, cyclin D, or cyclin E are common in solid tumors including breast cancers (Asghar et al., 2015; Balko et al., 2014). Also common are deficiencies in pRb function, which cause unregulated S phase entry, and deletion of the CDK4/6 inhibitor p16 (encoded by *CDKN2A*) (Asghar et al., 2015; Franco et al., 2014).

First generation pan-CDK inhibitors active against cell cycle regulators such as CDK1/2/4/6 and transcriptional regulators such as CDK9 arrest cells in both G1 and G2 and are broadly cytotoxic, and their clinical development has been challenged by poor therapeutic windows (Asghar et al., 2015). Subsequent generations of CDK inhibitors have been designed to inhibit specific CDK proteins (or subfamilies). In February 2015, the CDK4/6 inhibitor, palbociclib (PD0332991; Ibrance®) (Cristofanilli et al., 2016) received FDA approval for management of hormone receptor-positive (HR+) metastatic breast cancer (MBC) (Finn et al., 2009; O’Leary et al., 2016). Clinical trials of the CDK4/6 inhibitors, ribociclib (LEE011; KISQALI®) (Hortobagyi et al., 2016) and abemaciclib (LY2835219; Verzenio®) (Dickler et al., 2016; Sledge et al., 2017) also demonstrated substantial improvements in progression-free survival in HR+ metastatic breast cancer (Cristofanilli et al., 2016; Griggs and Wolff, 2017) leading to their approval by the FDA. CDK4/6 inhibitors are currently regarded as some of the most promising new drugs for the treatment of HR+ breast cancer and are also being tested against other malignancies (Goel et al., 2016; Lim et al., 2016; McCain, 2015; Patnaik et al., 2016a).

As observed with many other targeted therapies, acquired resistance to CDK4/6 inhibitors develops over time and nearly all initially responsive patients ultimately progress (Sherr et al., 2016). Resistance to CDK4/6 inhibitors is associated with multiple genomic alterations including amplification of Cyclin E, which promotes CDK2-dependent phosphorylation of pRb, amplification of CDK6, and loss of pRb function (Asghar et al., 2015; Yang et al., 2017). High expression of cyclin E is also associated with high CDK2 activity post-mitosis, which appears to bypass a requirement for CDK4/6 for cell cycle reentry (Asghar et al., 2017).

Despite having the same nominal targets and similar clinical indications, emerging evidence suggests that palbociclib, ribociclib, and abemaciclib differ in the clinic: abemaciclib in particular has been reported to have unique single-agent activities and distinct adverse effects (O’Brien et al., 2018; Patnaik et al., 2016b). The three drugs are dosed differently, have different pharmacokinetics, and are reported to differ with respect to target selectivity (Chen et al., 2016; Cousins et al., 2017; Gelbert et al., 2014; Kim et al., 2013). Among abemaciclib secondary targets examined to date, inhibition of DYRK/HIPK kinases is thought to contribute to cellular cytotoxicity (Knudsen et al., 2017); inhibition of GSK3α/β can activate WNT signaling (Cousins et al., 2017); inhibition of CDK9 is thought to be therapeutically unimportant (Torres-guzmán et al., 2017); however, overall the biological significance of differences in potency against primary CDK4/6 targets and secondary targets remains largely unexplored.

The target profiles of most clinical compounds are established relatively early in their development and are not necessarily revised at the time of approval. This is further complicated in the case of kinase inhibitors by the use of different measurement technologies to assess selectivity and the steady evolution of these technologies over the course of development of a single drug. By directly comparing the target profiles and biological activities of palbociclib, ribociclib and abemaciclib, as well as an earlier generation pan-CDK inhibitor, alvocidib (flavopiridol), we sought to address three related questions: (i) are the three approved CDK4/6 inhibitors interchangeable with respect to biochemical and cell-based activities; (ii) is there a possibility that tumors that have become resistant to one CDK4/6 inhibitor remain responsive to another inhibitor; and (iii) what are the relative merits of different approaches to characterizing the target spectra of kinase inhibitors?

In this paper we report the analysis of the clinically-approved CDK4/6 inhibitors using five experimental approaches that provide complementary insights into drug mechanisms of action: (i) mRNA sequencing of drug-perturbed cells, (ii) phosphoproteomics using mass spectrometry, (iii) GR-based dose-response measurement of cellular phenotypes (Hafner et al., 2016), (iv) mRNA sequencing of drug-treated xenograft tumors and (v) *in vitro* analysis for inhibitory activity using three different approaches: activity assays with recombinant enzymes; kinome-wide profiling using the commercial KINOMEscan platform from DiscoverX (Fabian et al., 2005); and kinase enrichment proteomics based on affinity purification on kinobeads (Duncan et al., 2012). We find that the five experimental approaches provide different but complementary views of target coverage and demonstrate that palbociclib, ribociclib, and abemaciclib have substantial differences in biological activities and secondary targets in breast cancer cell lines of varying genotypes. Multiple lines of evidence, including an *in vivo* xenograft model and preliminary data on patients and patient-derived cell lines treated with abemaciclib, suggest that the unique activities of abemaciclib arise from inhibition of kinases in addition to CDK4/6, notably CDK1 and CDK2 and may be therapeutically advantageous.

## RESULTS

### Approved CDK4/6 inhibitors induce distinct molecular signatures in breast cancer cells

To compare the mechanisms of action of palbociclib, ribociclib, and abemaciclib we performed transcriptional profiling (mRNA-seq) on a panel of seven breast cancer cell lines following 6 or 24 hours of exposure to 0.3, 1, or 3 μM of drug (Figure 1a and Table S1). In all but pRb-deficient BT-549 cells, treatment with any of the three drugs gave rise to a signature (signature 1; Figure 1a in red) comprising 87 significantly down-regulated genes (FDR < 0.2). In addition, treatment of cells with abemaciclib in the low micromolar range (gray box in Figures 1a) induced a second transcriptional signature (signature 2; Figure 1a in cyan) comprising 688 significantly down-regulated genes (FDR < 0.2) that was absent from ribociclib-exposed cells and only weakly present in cells exposed to palbociclib. We queried the Broad Connectivity Map (CMAP) (Lamb et al., 2006) with the two sets of down-regulated genes to determine which drug-induced changes they most closely matched. For signature 1, palbociclib and inhibitors of MEK (MAP kinase kinase) were the strongest hits (ribociclib and abemaciclib are absent from the CMAP dataset; Figure 1b and Table S2). Like CDK4/6 inhibition, MEK inhibition is anti-mitogenic in breast cancer cells, causing cells to arrest at the G1/S transition (Caunt et al., 2015; Meloche and Pouysségur, 2007). Gene set enrichment analysis showed that signature 1 was enriched for genes in the set *Reactome “Cell Cycle”* (p=9.0×10^−50^); it therefore appears to reflect cell cycle arrest in G1 (O’Leary et al., 2016). We scored the strength of this signature (the “G1-arrest score”) as the absolute mean log_2_ fold-change in the expression of all genes comprising signature 1. When signature 2 was compared to CMAP, the strongest hits were alvocidib and other pan-CDK inhibitors (Figure 1c and Table S2), suggesting that this signature arises from inhibition of CDKs other than CDK4 and CDK6. The strength of signature 2 (the “pan-CDK score”) was also calculated as the absolute mean log_2_ fold-change in gene expression. The G1-arrest score was high for all three drugs (Figure S1) whereas the strength of the pan-CDK score varied with drug and dose; it was highest for abemaciclib above 0.3 µM and lowest for ribociclib. Palbociclib exposure was associated with intermediate scores (Figure 1d).

**Figure 1:**
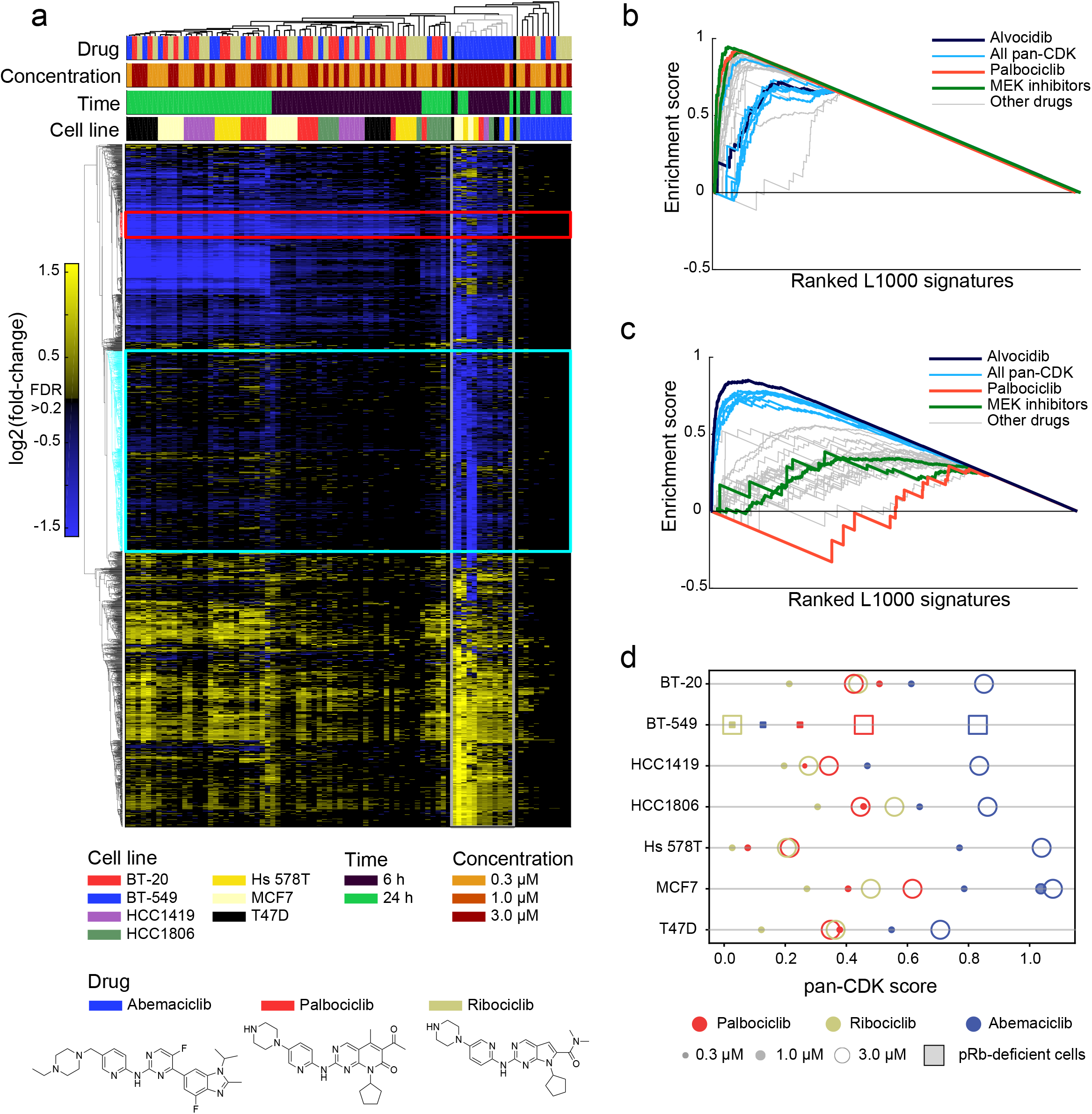
Transcriptional responses of breast cancer cell lines to CDK4/6 inhibitors. **(a)** Clustering of transcriptional responses for seven breast cancer cell lines treated for 6 or 24 hours with ribociclib, palbociclib, or abemaciclib at 0.3, 1, or 3 µM. Only genes for which statistically significant (FDR < 0.2) changes were observed in at least 3 conditions are shown. Down-regulated genes comprising signature 1 and 2 are outlined in red and cyan, respectively, and the gray box denotes the cluster containing expression profiles with the highest signature 2 scores. **(b-c)** Enrichment scores for signature 1 (b) and 2 (c) based on L1000 signatures identified by Enrichr (see Methods). **(d)** Score of the pan-CDK transcriptional signature per cell line following six hours of exposure to drug based on RNA-seq data from panel (a).

To better understand the origins of the pan-CDK signature, we collected RNAseq data from a larger set of conditions using the high-throughput, low-cost RNA sequencing method 3’ Digital Gene Expression (DGE-seq) (Soumillon et al., 2014). Seven cell lines, including two that are pRB-deficient (BT-549 and PDX-1258), were exposed for 6 hours to palbociclib, ribociclib, or abemaciclib or to alvocidib (which inhibits CDK1/2/4/6/9); data were collected in triplicate at four CDK4/6 inhibitor concentrations and two alvocidib concentrations. Differential expression of genes in signatures 1 and 2 (as defined above) was then used to compute G1-arrest and pan-CDK scores for each condition (Figure 2, Table S3). From these data we found that the strength of the average pan-CDK score was ordered as follows: alvocidib > abemaciclib > palbociclib > ribociclib (Figure 2, x-axis). For abemaciclib and alvocidib the pan-CDK score was strongly dose dependent (r=0.78, p=9.3×10^−7^ and r=0.76, p=1.5×10^−3^ respectively) as were the G1 arrest scores for all four drugs. Notably, the pan-CDK score for 0.1 µM and 1 µM alvocidib across all cell lines (green) substantially overlapped abemaciclib at 1 µM and 3 µM (blue).

**Figure 2:**
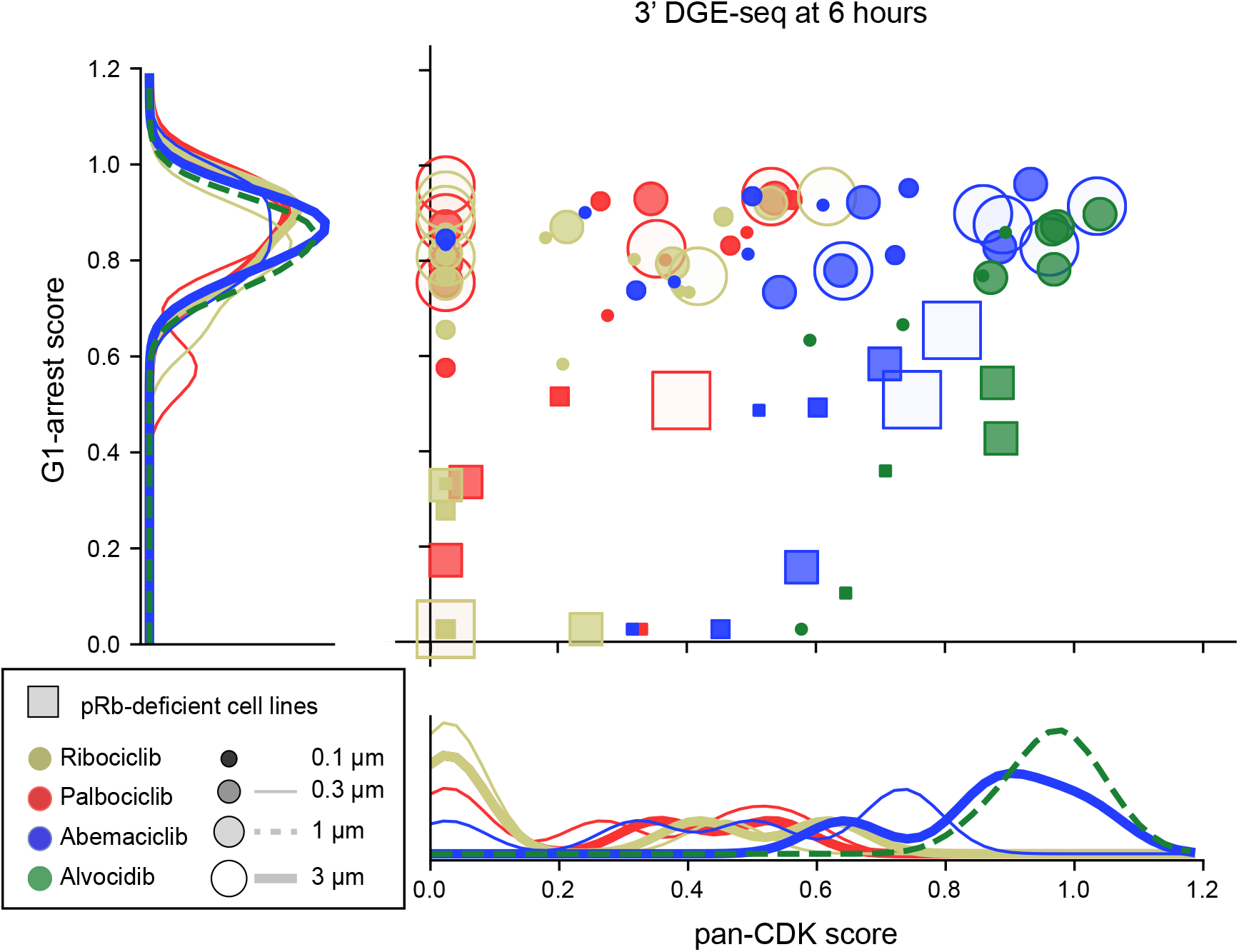
G1-arrest and pan-CDK scores induced by CDK4/6 inhibitors. Score of the G1-arrest signature relative to the pan-CDK signature for seven cell lines treated with palbociclib, ribociclib, abemaciclib, or alvociclib at 0.1, 0.3, 1, or 3 µM; squares denote pRb-deficient lines. Distributions of scores for pRb-competent lines are shown at the margins for each signature.

In pRb-deficient lines, only four genes in the G1 signature were differentially regulated by ribociclib (two-sided Fisher exact test p=2×10^−4^ as compared to pRb-proficient lines) consistent with the hypothesis that a pure CDK4/6 inhibitor should be inactive in cells lacking pRb, the primary substrate of CDK4/6 kinases. G1 arrest scores were lower in pRb-deficient than in pRb-proficient cell lines (0.25 vs. 0.73 on average) but they were not zero. This likely arises because pan-CDK and G1 arrest signatures are not orthogonal and inhibition of CDKs contributes to both. It was nonetheless true that high G1-arrest scores were observed when pan-CDK scores were near zero, particularly for ribociclib and palbociclib (data points lying along the y-axis in Figure 2). These RNA-seq data strongly suggest that palbociclib, ribociclib, and abemaciclib have different target spectra in breast cancer cells. Moreover, like alvocidib, abemaciclib is biologically active in pRb-deficient cells, as judged by changes in gene transcription.

### Effects of CDK4/6 inhibitors on the activity of CDK/cyclin complexes

To study the effects of CDK4/6 inhibitors on the phosphoproteome we performed isobaric (TMT) tag based liquid-chromatography mass spectrometry (LC/MS) (McAlister et al., 2012). MCF7 cells were treated with DMSO, palbociclib, or abemaciclib for one hour (to focus on immediate-early changes in the phosphoproteome) and a total of 9958 phosphopeptides were detected across all samples; among these phosphopeptides, 739 were down-regulated in the presence of palbociclib and 2287 in the presence of abemaciclib (log_2_ fold-change > 1.5; Figure 3a, Table S4). Enrichment analysis (Drake et al., 2012) involving known kinase-substrate relationships (see Methods) was used to infer changes in the activities of upstream kinases potentially accounting for observed changes in the phosphoproteome. The inferred activities for CDK4, CDK6, and Aurora A/B kinases (AURKA/B) were significantly lower in cells treated with either palbociclib or abemaciclib than a DMSO-only control whereas the activities of CDK1, CDK2, and CaM-kinase II subunit alpha (CAMKIIα) were lower only in cells treated with abemaciclib (Figure 3b, Table S5).

**Figure 3:**
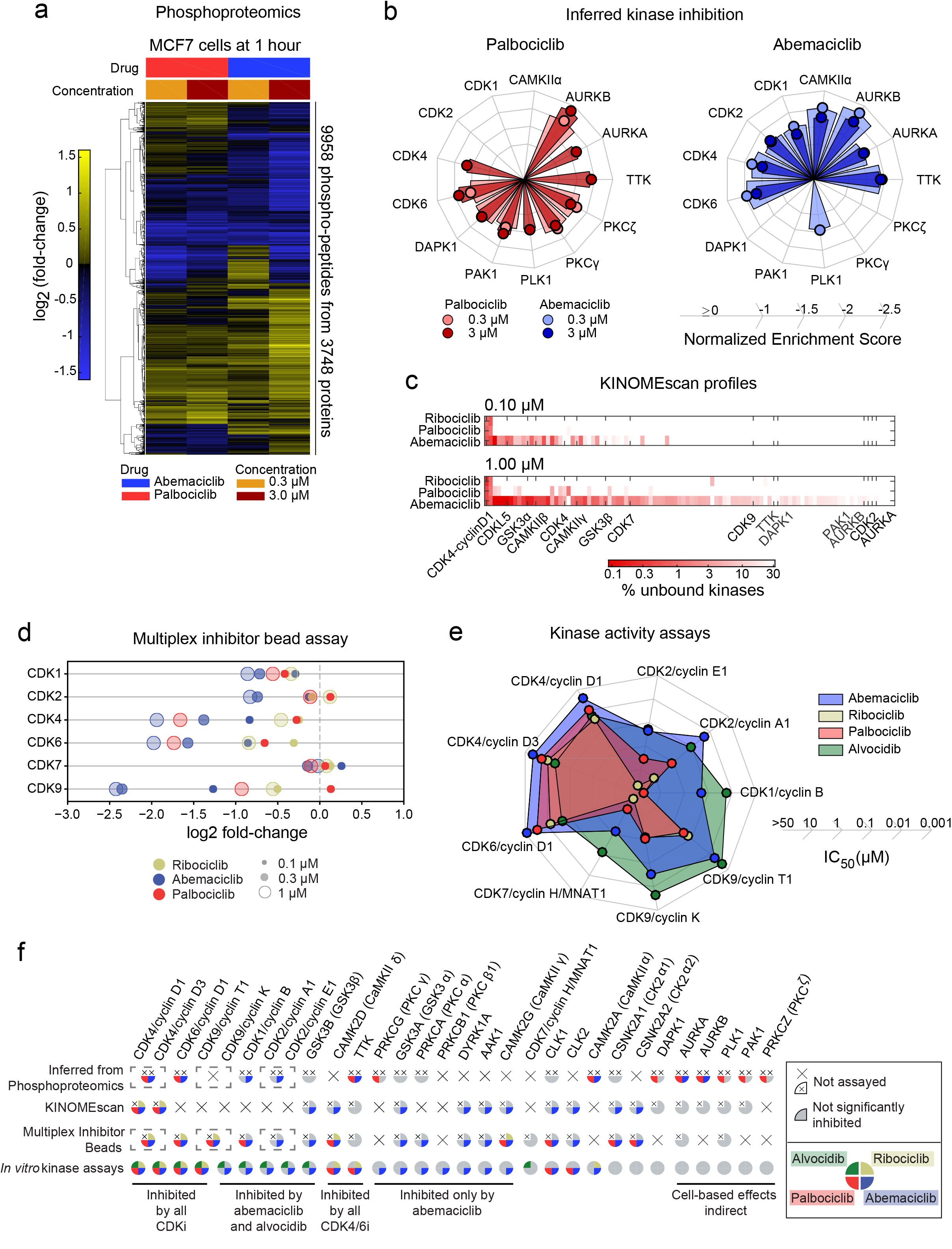
Inhibition of CDK/cyclin activity by CDK4/6 inhibitors. **(a)** Clustering of changes in phosphopeptide levels for MCF7 cells treated for 1 hour with either abemaciclib or palbociclib at 0.3 or 3 µM. **(b)** Normalized enrichment scores for kinases based on the phosphoproteomic data in panel (a). Only kinases inferred as significantly down-regulated (FDR < 0.2) in at least two conditions are shown. **(c)** Fraction of unbound kinases at 0.1 and 1 µM of each CDK4/6 inhibitor as measured by the KINOMEscan assay for the top 100 bound kinases plus kinases inferred in (b) (see Figure S2). CAMKIIα, CDK6, PKCγ, and PKCζ were not present in the panel. **(d)** Degree of inhibition (log_2_ fold change) of each CDK as detected by MIB/MS after treating a mixed cell lysate with a CDK4/6 inhibitor at the doses indicated. **(e)** *IC_50_* values for CDK/cyclin complexes for CDK4/6 inhibitors and alvocidib as measured using purified kinases *in vitro* (see Figure S3). **(f)** Summary of kinases that were assayed by phosphoproteomics, KINOMEscan, MIB/MS, and SelectScreen. Each slice of the pie represents inhibition by abemaciclib, ribociclib, palbociclib, or alvocidib. For each assay, slices are colored only if the corresponding drug substantially inhibited the kinases (defined as FDR <= 0.2 for phosphoproteomics inference, 90% inhibition at 1 µM drug by KINOMEscan, log_2_ fold-change < −0.45 for MIB/MS, or *IC*_50_ < 0.5 µM by in vitro kinase assays). An ‘x’ inside a slice denotes that that drug was not profiled in that assay. A large ‘X’ in place of a pie indicates that that kinase was not profiled in that assay. Bound complexes such as CDK4/cyclin D1or CDK4/cyclin D3 cannot be disambiguated in kinase inference and MIB/MS assays and are therefore depicted as a single entity within a box.

Kinase inference suggests that palbociclib and abemaciclib down-regulate the activities of multiple kinases other than CDK4 and CDK6. However, this conclusion has several caveats, most importantly, that kinase inhibitors can act indirectly, for example, by blocking the activity of an upstream kinase in a multi-step cascade or by arresting cells at a point in the cell cycle at which some kinases are not normally active (CDKs for example). Second, even when using state-of the-art mass spectrometers and methods, less than 10% of the total phosphoproteome can be analyzed in any single sample, making kinase-inference subject to statistical error. Third, there exists a poorly established many-to-many mapping between kinases and substrates, necessitating predictive models based on motif signatures and binding probabilities, all with associated uncertainties. Because of these limitations we consider kinase inference to be a semi-quantitative method: large differences across drugs are likely meaningful but dose-response relationships can be hard to capture.

To distinguish direct and indirect effects of kinase inhibitors on the phosphoproteome we performed three different type *in vitro* assays. First, we used the commercial KINOMEscan assay, which measures binding between members of a 468 DNA-tagged recombinant kinase library and an immobilized ATP-like ligand; the assay is performed in the presence and absence of an ATP-competitive drug (Fabian et al., 2005). KINOMEscan profiling showed that ribociclib is the most selective CDK4/6 inhibitor and abemaciclib the least (Figure 3c, Figure S2a-b and Table S6). KINOMEscan assays have previously been performed on CDK4/6 inhibitors (Chen et al., 2016; Gelbert et al., 2014); our data agree with earlier findings.

Several CDKs are not found in the KINOMEscan library (e.g. CDK1, CDK6) or are not complexed with cyclins (e.g. CDK2); therefore, we used a second method to obtain kinome profiles: multiplexed inhibitor bead mass spectrometry (MIB/MS) (Duncan et al., 2012). In this approach, a cell lysate is mixed with beads conjugated to pan-kinase inhibitors in the presence and absence of a test drug and the levels of bound kinases then determined by mass spectrometry (Figure 3d, Table S7); to generate a lysate with the greatest number of expressed kinases, we mixed several cell types (Médard et al., 2015). We detected 164 kinases, including 13 CDKs in the unfractionated extract by TMT LC/MS, and found that ribociclib, palbociclib, and abemaciclib all bound to CDK4 and CDK6. In addition, abemaciclib bound to CDK1, CDK2, CDK7, CDK9, GSK3α/β and CAMKIIγ/δ. These results agree well with data for abemaciclib recently published by Cousins et al. (Spearman’s ρ = 0.62, *P* = 8.9×10^−16^) (Cousins et al., 2017). Moreover, when KINOMEscan data (obtained in the presence of 1 µM abemaciclib) and MIB data (obtained with 10 µM abemaciclib) were compared, 19 of 25 kinases strongly inhibited in the KINOMEscan and also present in cell extracts were significantly reduced in binding to MIBs (log2 fold change > 1.5), demonstrating good reproducibility between different types of assays. We conclude that ribociclib is the most selective CDK4/6 inhibitor tested and abemaciclib the least, with a dose-dependent increase in the number of targets significantly inhibited by abemaciclib from 4 at 0.1 µM drug to 13 at 1 µM and 28 at 10 µM.

As a third approach, we performed *in vitro* kinase activity assays at 10 concentrations (using SelectScreen technology by Thermo Fisher and HotSpot technology by Reaction Biology; see Methods). Drugs were tested on the kinases and kinase-cyclin complexes that we identified as potential abemaciclib targets by transcriptional, phospho-proteomic or kinase profiling assays. The data showed that abemaciclib was the most potent inhibitor of CDK4 and CDK6 of the three drugs tested and that it was also active against multiple kinases that were not inhibited, or were only weakly inhibited, by palbociclib or ribociclib (Figure 3e, Figure S3 and Table S8). These kinases include CDK2/cyclin A/E, CDK1/cyclin B, CDK7/cyclin H, CDK9/cyclin K/T1, CAMKIIα/β/γ, and GSK-3α/β (Figure 3e, Figure S3). Compared to the first-generation CDK inhibitor alvocidib, abemaciclib had similar potency against CDK2/cyclin A/E but was ~10-fold less potent against CDK1/cyclin B, CDK7/cyclin H, and CDK9/cyclin K/T1 (potentially explaining the improved toxicity profile of abemaciclib relative to pan-CDK inhibitors), whereas ribociclib and palbociclib were at least another order of magnitude less potent than abemaciclib against these secondary targets. The potency of the three drugs against CDK4 vs. CDK6 was dependent on the cyclin partner and the assay, but the kinases generally differed by no more than 3-fold (Table S8).

Results from KINOMEscan, MIB/MS, and SelectScreen assays performed *in vitro* were largely concordant with mRNA-seq and phosphoproteome profiling with a few notable exceptions (Figure 3f). CDK1 and CDK6 were absent from the KINOMEscan panel and CDK2 was not found to be a target, probably because the appropriate cyclin was absent and cyclin binding changes CDK2 activity (Echalier et al., 2014). Such a false-negative result in the widely used KINOMEscan assay may explain why the activity of abemaciclib against CDK2-cyclin A/E is under-appreciated. Biochemical assays showed that abemaciclib was inactive against other kinases such as AURKA/B, and PAK1 (Figure S3a) and the downregulation inferred from phosphoproteomic data most likely reflects an indirect effect: arrest of cells in G1 by CDK4/6 inhibition is expected to block normal phosphorylation of AURKA/B, and PAK1 in G2/M phase. Thus, it was only by combining multiple *in vitro* and cell-based assays that a complete picture of kinase inhibitor activities was obtained (Figure 3f).

### Comparing CDK4/6 inhibitors in breast cancer cell lines

To compare the biological activities of CDK4/6 inhibitors, we acquired dose-response curves in 34 breast cancer cell lines spanning all clinical subtypes and computed GR values (Figure 4a and Table S9) which distinguish between drug-induced cell cycle arrest and cell death while correcting for artifactual differences in drug sensitivity arising from variability in proliferation rates (Hafner et al., 2016, 2017). Both palbociclib and abemaciclib elicited cytostatic responses with *GR_50_* values in the 10-100 nM range (Table S10). Potency was highly correlated between the drugs (Spearman’s ρ = 0.91, *P* = 5.7×10^−14^) with abemaciclib ~5.5-fold more potent on average at inducing cytostasis (t-test *P* = 5.3×10^−7^); this difference is consistent with a 3-fold difference between palbociclib and abemaciclib in *in vitro IC_50_* values for CDK4/6 kinase activity (Figure 3e). Efficacy at 0.1 µM drug, as measured by GR value, varied between 0 (complete cytostasis) and 0.76 (weak growth inhibition) in pRb-proficient cell lines but was similar for palbociclib and abemaciclib, showing that at these concentrations the drugs induce similar phenotypic effects and only fractionally inhibit cell proliferation. In pRb-deficient cell lines, palbociclib was inactive at all doses and abemaciclib had little or no effect below 0.3 µM (yellow lines Figure 4a). The cytostatic response observed at lower abemaciclib doses and all doses of palbociclib is most likely a result of CDK4/6 inhibition.

**Figure 4:**
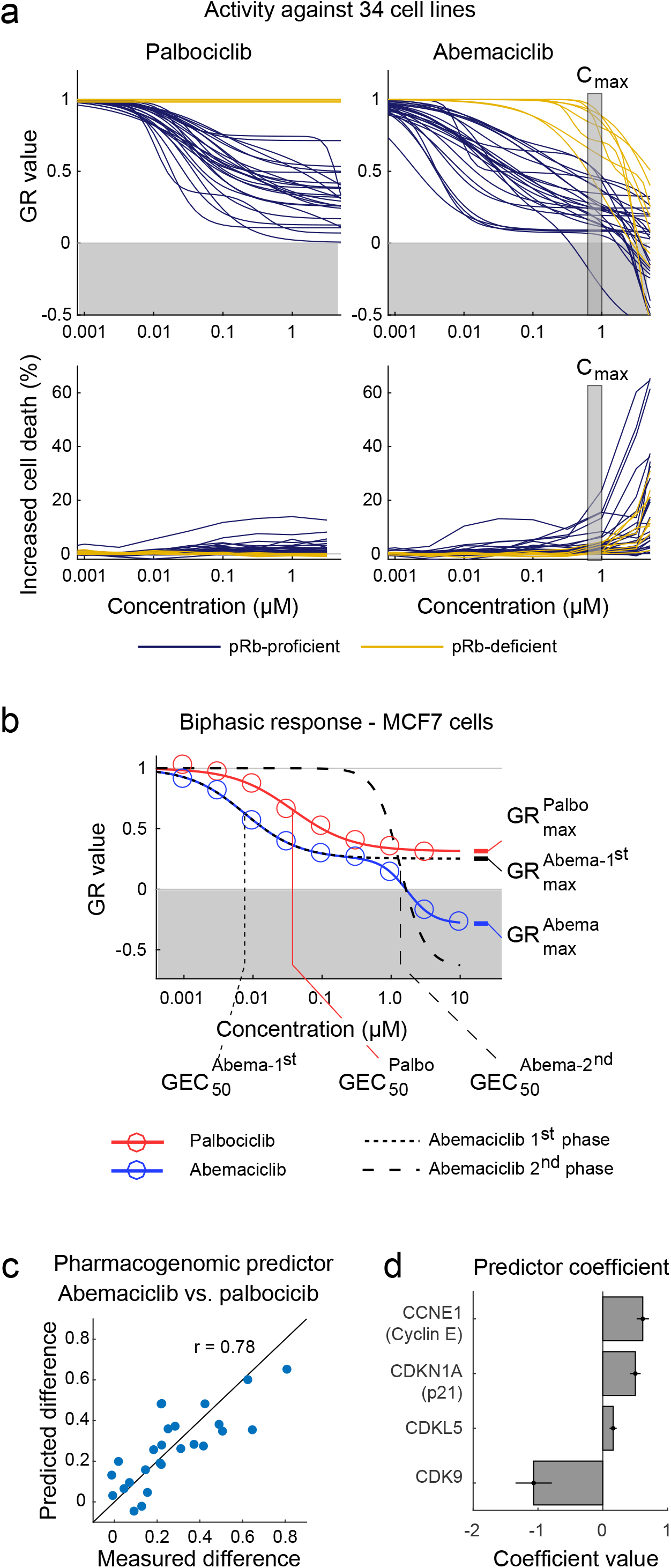
Comparison of the phenotypic response of breast cancer cell lines to CDK4/6 inhibitors. (**a**) GR curves for cell growth (top) and increase of dead cells relative to a vehicle-only control (bottom) for 26 pRb-proficient breast cancer cell lines (blue) and 8 pRb-deficient cell lines (yellow) treated with palbociclib (left) or abemaciclib (right) for 72 hours. The vertical box illustrates the maximum serum concentration for abemaciclib (*C_max_*). (**b**) Dose-response curve for palbociclib (red) and abemaciclib (blue) in MCF7 cells. Dotted lines depict two fitted sigmoidal curves whose product optimally recapitulates the blue curve with extracted values for *GEC_50_* (50%-maximal effective concentration) shown below and for *GR_max_* (maximal efficacy) shown to the right (See Figure S4). (**c-d)** Performance of a pharmacogenomic predictor of palbociclib vs. abemaciclib drug response constructed from data on mRNA levels for 30 cell cycle regulators; (**c**) shows the observed versus predicted (leave-one-out cross validation) difference in GR value at 3 µM between palbociclib and abemaciclib based on a linear model containing the expression of four genes, whose coefficients are shown in (**d**); error bars represent the standard error of the model.

However, abemaciclib also elicited a second response at doses greater than 0.3 µM; this response was characterized by negative GR values and cell death (see Methods; Figure 4a). As a result, the complete dose-response behavior of abemaciclib was significantly better fitted in most cell lines by the product of two sigmoidal curves (Figure 4b, Figure S4, and Methods). The mid-point of the second response curve was offset to a similar degree as *in vitro* dose-response curves for CDK1/2 vs. CDK4/6 inhibition (Table S8). This behavior is consistent with inhibition of two sets of targets: CDK4/6 at low dose – resulting in G1 arrest – and kinases such as CDK1/2 above 0.3 µM – resulting in cell death. At all doses tested in all cell lines, responses to palbociclib and ribociclib were purely cytostatic (GR > 0). As a result, abemaciclib was substantially more efficacious than palbociclib in inhibiting and killing pRb-proficient cells of all subtypes, having a *GR_max_* value on average 0.52 below that of palbociclib (t-test *P*=4.5×10^−9^; Table S10).

A search of 30 cell cycle regulators for genes whose mRNA expression levels could discriminate between responsiveness to 3 µM palbociclib and abemaciclib in the 26 pRb-proficient cell lines yielded a high-performing multi-linear model involving only four genes (*q^2^* = 0.85, *P* = 2.9×10^−6^ by leave-one-out cross validation; Figures 4c-d and S4). The genes were CCNE1 (cyclin E1), which has been implicated in palbociclib resistance (Sherr et al., 2016; Turner et al., 2019), CDKN1A (p21 – an inhibitor of CDK1/2/4/6), CDK9 (a target of abemaciclib and pan-CDK inhibitors), and CDKL5 (cyclin-dependent kinase-like 5). Our data showed CDKL5 to be strongly inhibited by abemaciclib (*IC_50_* ~ 18 nM in vitro) but not by palbociclib or ribociclib (*IC_50_*>3 µM or >10 µM respectively; Table S8). Thus, differences in the efficacy of CDK4/6 inhibitors on cell lines are related to the expression level of genes targeted uniquely by abemaciclib.

### Abemaciclib blocks cells in the G2 phase of the cell cycle

Consistent with the known biology of CDK4/6 inhibition, abemaciclib, ribociclib, and palbociclib all suppressed pRb phosphorylation and arrested cells in G1 (Figure 5). The 3-fold difference in drug concentration needed to induce arrest matched measured differences in potency in biochemical assays (with abemaciclib the most potent and ribociclib the least; Figure 3e). A fraction of cells treated with abemaciclib also arrested in G2 rather than G1, particularly at drug concentrations of 0.3 µM and above (Figure 5, Figure S5), a possible consequence of inhibition of CDK1 and CDK2, whose activities are required for progression through S-phase and mitosis. Treating pRb-deficient cells with ribociclib or palbociclib had no effect on cell cycle distribution whereas treatment with abemaciclib caused cells to accumulate in G2, consistent with an abemaciclib-induced cell cycle arrest independent of CDK4/6 (Figure 5, Figure S5).

**Figure 5:**
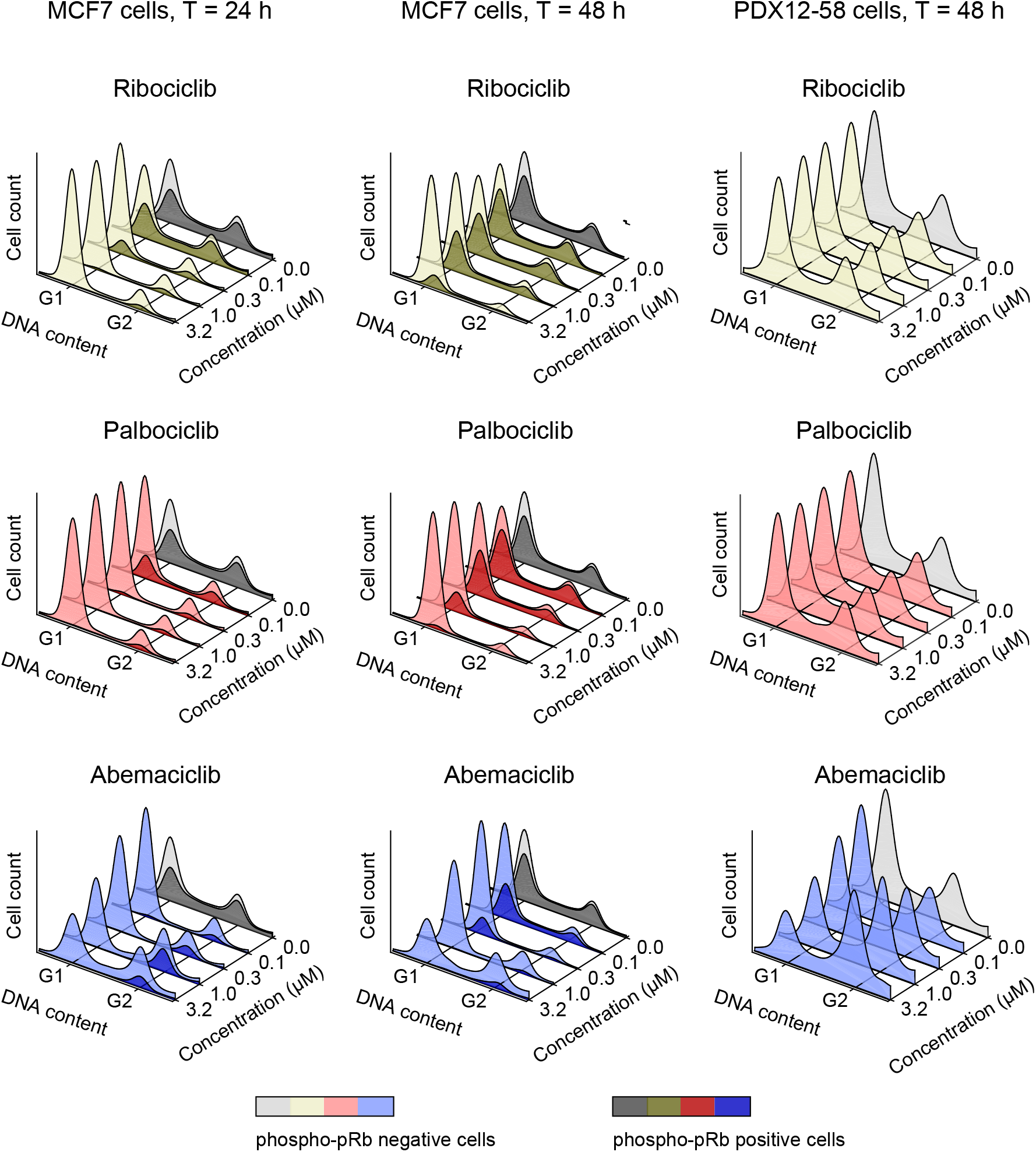
Comparison of the effects of ribociclib, palbociclib, and abemaciclib on the cell cycle. Distribution of DNA content in MCF7 cells exposed to one of three CDK4/6 inhibitors over a range of concentrations for 24 (left) or 48 (middle) hours, and in PDX-1258 cells, which are pRb-deficient, exposed to the same conditions for 48 hours (right). In each curve the phospho-pRb positive cell population is depicted in a darker shade. One representative replicate out of three is shown.

### Assaying abemaciclib polypharmacology in xenograft tumors

When a drug inhibits multiple targets with different potencies the question arises whether both primary and secondary targets can be engaged at doses achievable *in vivo*. When we compared G1-arrest and pan-CDK signature scores and cellular phenotypes across a range of abemaciclib doses in multiple cell lines, we found that pan-CDK scores were significant only above 0.3 µM (*P*=2.1×10^−4^, ranksum test) and cytotoxicity was observed in pRb-deficient cells only at concentrations of 1 µM and above. This compares well with a maximum serum concentration in humans (*C_max_*) for abemaciclib of 0.5 µM to 1 µM when active metabolites are included (Burke et al., 2016; Patnaik et al., 2016a). As a direct test of *in vivo* activity we generated MCF-7 xenografts in nude mice and exposed them to CDK4/6 inhibitors at a range of doses. When tumors reached ~300 mm^3^, animals were randomly assigned to treatment groups and treated daily for 4 days to a vehicle-only control or to 150 mg/kg ribociclib, 150 mg/kg palbociclib or 25-150 mg/kg abemaciclib, doses previously shown to be effective in xenografts (Fry et al., 2004; Gelbert et al., 2014; O’Brien et al., 2014). Animals were euthanized and tumors divided into two samples; one was fixed in formaldehyde and paraffin embedded and the other processed for mRNA-sequencing. FFPE specimens were imaged by immunofluorescence using vimentin and E-cadherin staining to distinguish tumor cells from mouse stroma.

We found that all conditions tested resulted in a significant reduction in the fraction of p-pRb positive cells (Dunnett’s multiple comparison *P* < 0.0001) providing pharmacodynamic evidence that all tumors were exposed to drug at active concentrations (Figure 6a). mRNA-seq data showed that all three drugs induced a G1-arrest signature (Figure 6b, Table S11), the strength of which was correlated with the degree of p-pRb inhibition (Spearman’s ρ = −0.80, *P* = 1.1×10^−10^). Furthermore, at doses above 100 mg/kg, abemaciclib (but not ribociclib or palbociclib) also induced a strong pan-CDK signature (Figure 6b). These data provide *in vivo* confirmation that abemaciclib can engage targets other than CDK4 and CDK6, recapitulating data on the drug’s off-target activity in cell culture.

**Figure 6:**
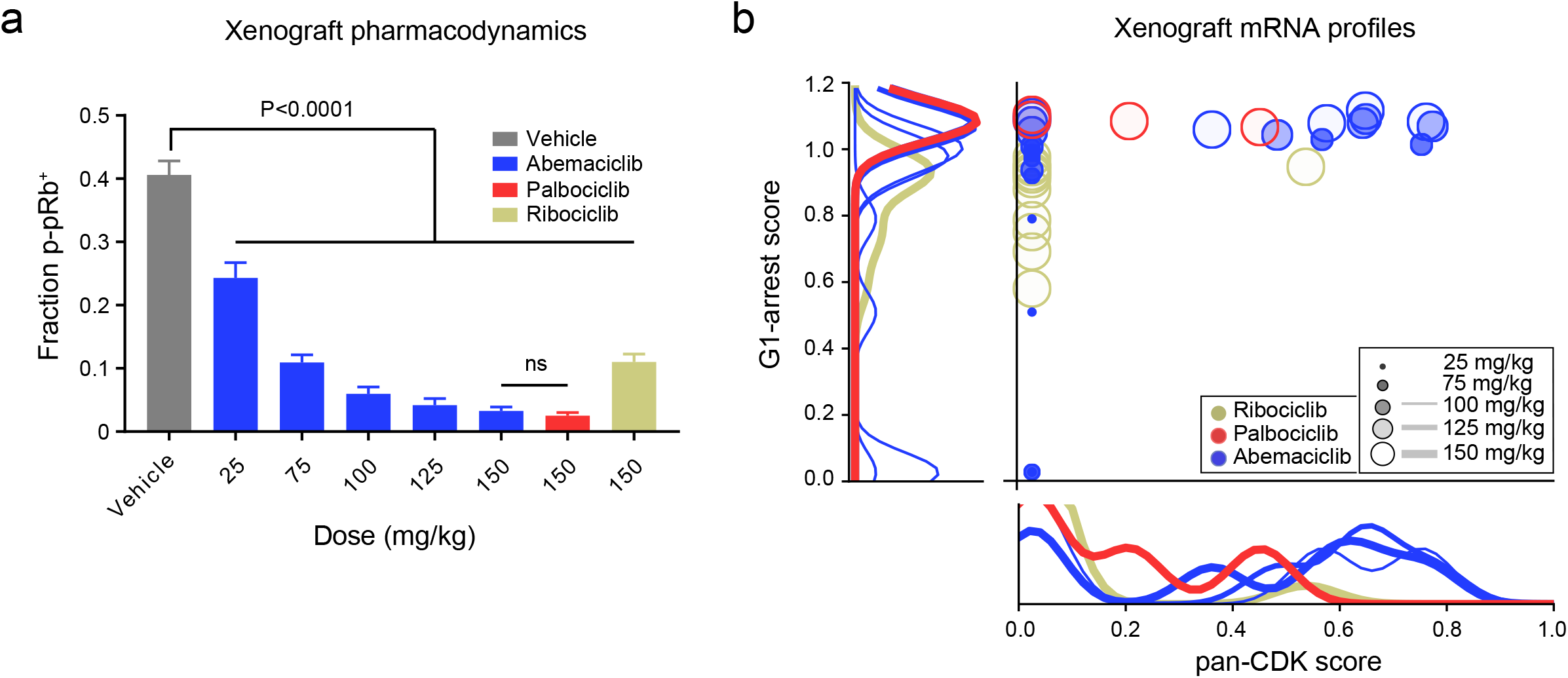
Transcriptional response of MCF-7 xenografted cells to CDK4/6 inhibitors. (**a**) Fraction of phospho-pRb positive tumor cells in MCF-7 xenografts after four days of CDK4/6 inhibitor treatment. (**b**) Score of the pan-CDK transcriptional signature as compared to the G1-arrest signature across MCF-7 tumors following four days of exposure to drug; same analysis as in Figure 2.

### Cross-resistance between abemaciclib and palbociclib or ribociclib is incomplete

As previously described (Asghar et al., 2017; Herrera-Abreu et al., 2016), cells adapt to CDK4/6 inhibition over time. Within 48 hours of exposure to palbociclib or ribociclib we found that cells re-entered the cell cycle and acquired a p-pRb positive state at drug concentrations as high as 3.16 µM (Figure 5a). In contrast, pRb phosphorylation remained low in cells exposed to 1 µM or more abemaciclib (Figure 5a) with ongoing cell death and no evidence of adaptation five days after drug exposure (Figure 7a, Figure S6 and Table S12). In studies designed to assess long-term adaptation to drug, we observed that breast cancer cells grown for several months in the presence of 1 µM palbociclib had higher cyclin E (CCNE1) and lower pRb levels than parental cells (Figure 7b). These palbociclib-adapted cells were cross-resistant to ribociclib (Figure 7c, Figure S7a-b and Table S13) but sensitive to abemaciclib at doses of 1 µM and above, consistent with the ability of abemaciclib to target kinases not inhibited by palbociclib.

**Figure 7:**
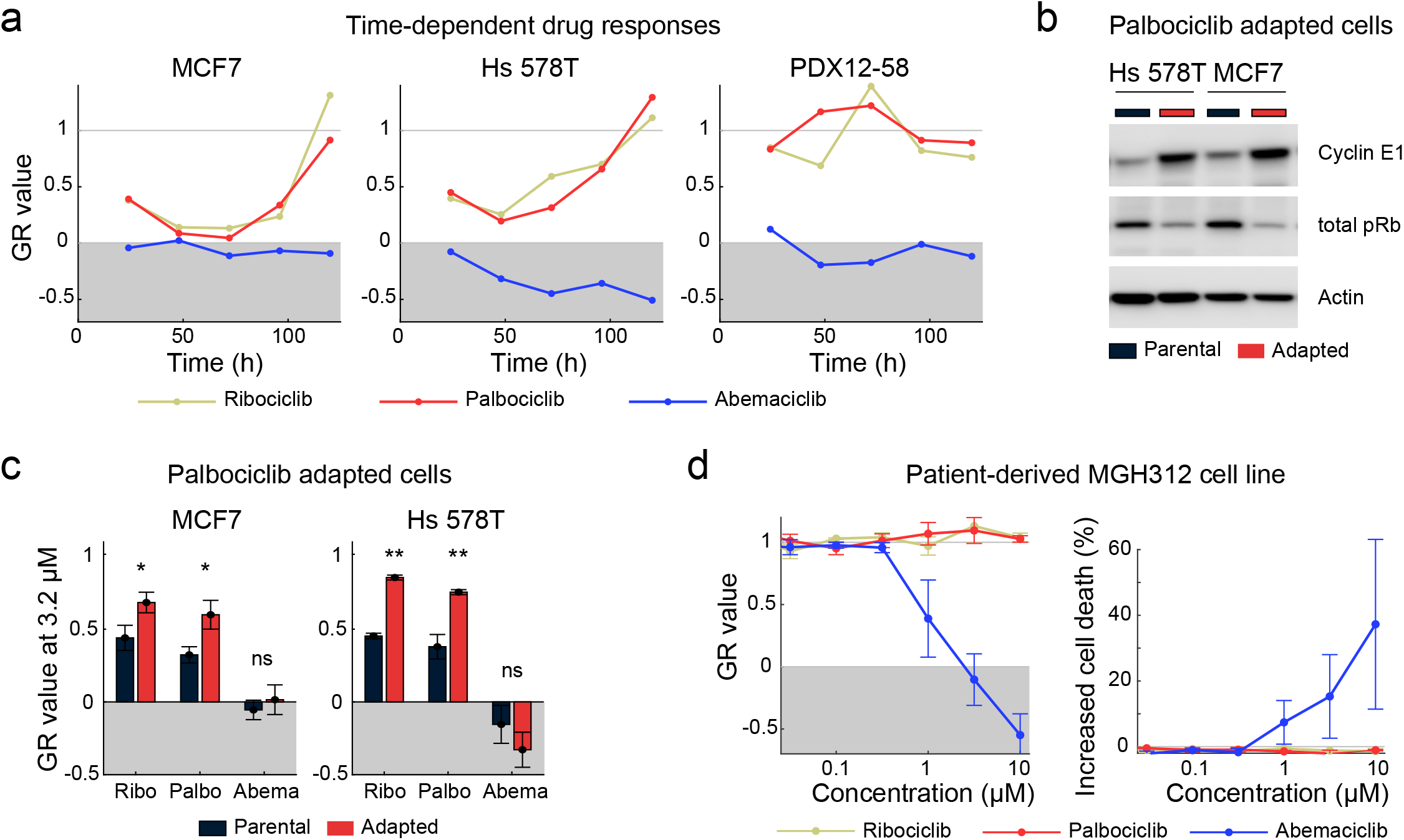
Acute and adaptive responses of breast cancer cell lines and tumors to CDK4/6 inhibitors. **(a)** Time-dependent GR values for MCF7, Hs 578T, and PDX12-58 cells treated with 3.16 μM ribociclib, palbociclib, or abemaciclib for up to five days. One representative replicate out of four is shown. **(b)** Western Blots of cyclin E and total pRb levels in Hs 578T and MCF7 parental cells and in cells adapted to grow in 1 µM palbociclib. **(c)** GR values for Hs 578T and MCF7 parental cells and cells adapted to grow in 1 µM palbociclib following exposure to 3.16 µM ribociclib, palbociclib, or abemaciclib for 72 h (see Figure S7); * denotes *P* < 0.05 and ** *P* < 0.01 as measured using a t-test with six replicates in each group. Error bars denote SEM of six replicates. **(d)** GR values (left) and increase in dead cells relative to a vehicle-only control (right) for the patient-derived line MGH312 in response to 96-hour exposure to ribociclib, palbociclib, or abemaciclib. Error bars show the SEM of three replicates.

We observed similar differences in a cell line established from a patient with advanced/metastatic HR+/Her2-breast cancer whose disease had progressed following eight months on ribociclib/letrozole. The tumor lacked pRb by immunohistochemistry (Figure S7c) as did the derived cell line (MGH312; Figure S7d). The tumor cells were responsive to abemaciclib as judged by inhibition of cell proliferation and induction of cell death but were completely resistant to palbociclib or ribociclib even at high doses (Figure 7d and Table S14). The potential for abemaciclib to benefit this patient remains unknown because she was now deceased and was never treated with abemaciclib.

## DISCUSSION

It is not uncommon for multiple therapeutics targeting the same proteins to be approved in close succession. In the case of CDK4/6 inhibitors, palbociclib, ribociclib and abemaciclib have all proven highly effective in the treatment of HR+ metastatic breast cancer and are currently being tested in ~100 ongoing clinical trials for activity in other malignancies. It has hitherto been assumed that the mechanisms of action of the three drugs are very similar, and distinct from those of older-generation CDK inhibitors such as alvocidib: observed differences in the efficacy and toxicity of palbociclib, ribociclib and abemaciclib have generally been attributed to differences in dosing schedules or potency against CDK4 versus CDK6 (Sherr et al., 2016). However, our work presents six lines of evidence that alvocidib, abemaciclib, palbociclib and ribociclib actually span a spectrum of increasing selectivity for CDK4/6-cyclin complexes. In particular, abemaciclib has biochemical and physiological activities not manifest by ribociclib and only weakly by palbociclib.

First, exposure of breast cancer cells of different subtypes to any of the three approved CDK4/6 inhibitors induces transcriptional changes associated with G1 arrest but abemaciclib alone induces dose-dependent transcriptional changes similar to those elicited by alvocidib reflective of pan-CDK inhibition. Second, exposing cells to abemaciclib results in more extensive changes in the phosphoproteome than exposure to palbociclib and kinase inference suggests that this is due in part to inhibition of CDK1 and CDK2. Third, kinome profiling using industry-standard KINOMEscan panels, multiplexed inhibitor bead mass spectrometry, and kinase activity assays confirms that abemaciclib has multiple targets in addition to CDK4/6. Fourth, abemaciclib causes arrest of cells in both the G1 and G2 phases of the cell cycle and the drug is cytotoxic (at high concentrations) even in the absence of pRb; in contrast, cells exposed to palbociclib and ribociclib arrest only in G1 and elicit little or no cell death. The difference in efficacy between abemaciclib and other CDK4/6 inhibitors is greatest in cell lines with a specific transcriptional profile (high CCNE1, CDKN1A and CDKL5 and low CDK9 expression levels). Fifth, in a mouse xenograft model, abemaciclib induces both CDK4/6-like G1 arrest and pan-CDK transcriptional signatures, as observed in cultured cells. Sixth, whereas abemaciclib durably inhibits cell division, cultured cells adapt within 2-3 days of continuous exposure to palbociclib or ribociclib and resume proliferation. Preliminary evidence of the clinical significance of these findings is provided by an abemaciclib-sensitive, palbociclib- and ribociclib-resistant cell line from a deceased patient with HR+/Her2-breast cancer who had progressed on ribociclib/letrozole.

Evidence of substantial differences among CDK4/6 inhibitors is scattered throughout the literature but has not been consolidated or rigorously evaluated, consistent with a general lack of comparative biochemical data on many FDA-approved drugs. Large-scale kinase profiling studies using KINOMEscan or MIB/MS are one exception to this generalization (Cousins et al., 2017; Fabian et al., 2005; Gelbert et al., 2014; Klaeger et al., 2017). However, our findings strongly argue for a multi-faceted approach to comparative mechanism of action studies. Proteomic, transcriptional, biochemical, and phenotypic approaches measure different aspects of drug action and, in the current work, a combination of methods was needed to obtain an accurate and complete picture of target spectrum. For example, the false negative finding in KINOMEscan data that abemaciclib does not interact with CDK2 may explain why biological differences among CDK4/6 inhibitors have not been widely appreciated. Similarly, whereas GSK3β was found to be an abemaciclib target of borderline significance by phosphoproteome profiling (perhaps as a result of proteome under-sampling (Riley and Coon, 2016)), it was clearly a target by kinase activity assays (Cousins et al., 2017). Conversely, proteomic profiling assays suggesting that abemaciclib exposure results in downregulation of AURKA/B and PLK1 activities is most likely an indirect consequence of cell cycle arrest. In agreement with Cousins et al. (Cousins et al., 2017), our results using multiple different assays provide little support for the assertion that ribociclib, palbociclib or abemaciclib are systematically more active against CDK4 than CDK6 (Gelbert et al., 2014; Patnaik et al., 2016a, 2016b). Although, enzymatic assays show the *IC_50_* for CDK4 is about 2.5-fold greater than for CDK6 for all three drugs, this is unlikely to be therapeutically significant because both targets are strongly inhibited at doses used in patients.

In the case of a polyselective drug such as abemaciclib the question arises whether activities observed at different drug concentrations are all biologically relevant. There is no question that CDK4 and CDK6 are the highest affinity targets of abemaciclib and that abemaciclib is the most potent of the three approved drugs against these CDKs. Our data show abemaciclib to be 10- to 100-fold less potent against CDK2 and CDK1 than CDK4/6, but we detect the cellular consequences of CDK1/2 inhibition in cell lines at concentrations as low as 0.3 µM, well within the *C_max_* range in humans, and also achievable in xenograft mouse models (Burke et al., 2016; Patnaik et al., 2016a; Raub et al., 2015). Abemaciclib also exhibits substantially reduced drug adaptation with respect to anti-proliferative effects, which is beneficial for an anti-cancer drug.

The current generation of CDK4/6 inhibitors has benefited from a considerable investment in increasing target selectivity, mainly as a means of reducing toxicity relative to earlier generation drugs (Asghar et al., 2015; Peplow, 2017; Toogood et al., 2005). However, our findings suggest that abemaciclib is not equivalent to palbociclib or ribociclib. Its activities against kinases other than CDK4/6 may be beneficial for anti-cancer activity and targeting them jointly with CDK4/6 may be a means to achieve more durable responses than with CDK4/6 inhibition alone. Inhibition of CDK1/7/9 may also contribute to cell killing (Kitada et al., 2000; Wittmann et al., 2003) and inhibition of mitotic kinases such as TTK may enhance tumor immunogenicity, a key contributor to drug response (Luen et al., 2016). Blocking CDK2/cyclin E should mitigate resistance resulting from amplification of cyclin E (a resistance mechanism in cell culture (Dean et al., 2010; Herrera-Abreu et al., 2016)) and also achieve a more complete therapeutic response by targeting mitotic cells with high CDK2 activity (Asghar et al., 2017). Patients whose tumors exhibit high expression levels of cyclin E1 and are non-responsive to palbociclib (Turner et al., 2019) may represent another cohort who might benefit from abemaciclib.

## SIGNIFICANCE

The integration of multiple cell-based and *in vitro* profiling methods has made it possible to directly compare the target spectra of three recently approved CDK4/6 inhibitors regarded as breakthroughs in the treatment of HR+ breast cancer. The substantially wider spectrum of activities detected for abemaciclib relative to other CDK4/6 inhibitors provides a rationale for treating patients with abemaciclib following disease progression on palbociclib or ribociclib. Cells, including cells derived from a breast cancer patient treated with ribociclib plus letrozole, who acquired resistance to these two drugs remain sensitive to abemaciclib at concentrations of 0.3 µM and above, overlapping human C*_max_* concentrations. The possibilities for use of abemaciclib in tumors that are pRb-deficient remain less certain, since drug activity is observed only at micromolar concentrations in pRb-deficient cell lines. A final possibility suggested by this work is combining CDK4/6 inhibitors with drugs that inhibit secondary targets of abemaciclib such as CDK2, a strategy Pfizer is pursuing in a single molecule (US patent 20180044344A1). In all of these cases, our work shows that polypharmacology can be exploited to achieve more durable responses than with “pure” CDK4/6 inhibitors such as ribociclib. More generally, our data suggest the value of systematic comparative target profiling of human therapeutics developed against the same targets but having different chemical structures.

## Acknowledgements

This work was funded by P50-GM107618, U54-CA225088 and U54-HL127365 to PKS and DJ. We thank LSP member M. Berberich for skilled assistance, the ICCB for help with automation, S. Gygi assistance with proteomics and A. Bardia for comments.

## Author contributions

MH, CEM, and DJ conceived the study; MH, CEM, DJ, and PKS designed the experiments and CEM, MC, SAB, and RAE performed them. MH, KS, and CC performed the computational analyses. CSW and DJ obtained the patient-derived line and provided related data. PKS oversaw the experimental and computational research; MH, CEM, KS, DJ, and PKS wrote the manuscript.

## Declaration of interests

MH is currently an employee of Genentech, Inc and RAE of Pfizer, Inc; they declare no conflicts of interest. DJ reports personal fees from Novartis, Genentech, Eisai, Ipsen, and EMD Serono, during the conduct of the study. PKS is a member of the SAB or Board of Directors of Merrimack Pharmaceutical, Glencoe Software, Applied Biomath and RareCyte Inc, has equity in these companies, and declares that none of these relationships are directly or indirectly related to the content of this manuscript. Other authors have no conflicts of interest.

## SUPPLEMENTAL FIGURE LEGENDS

**Figure S1.**
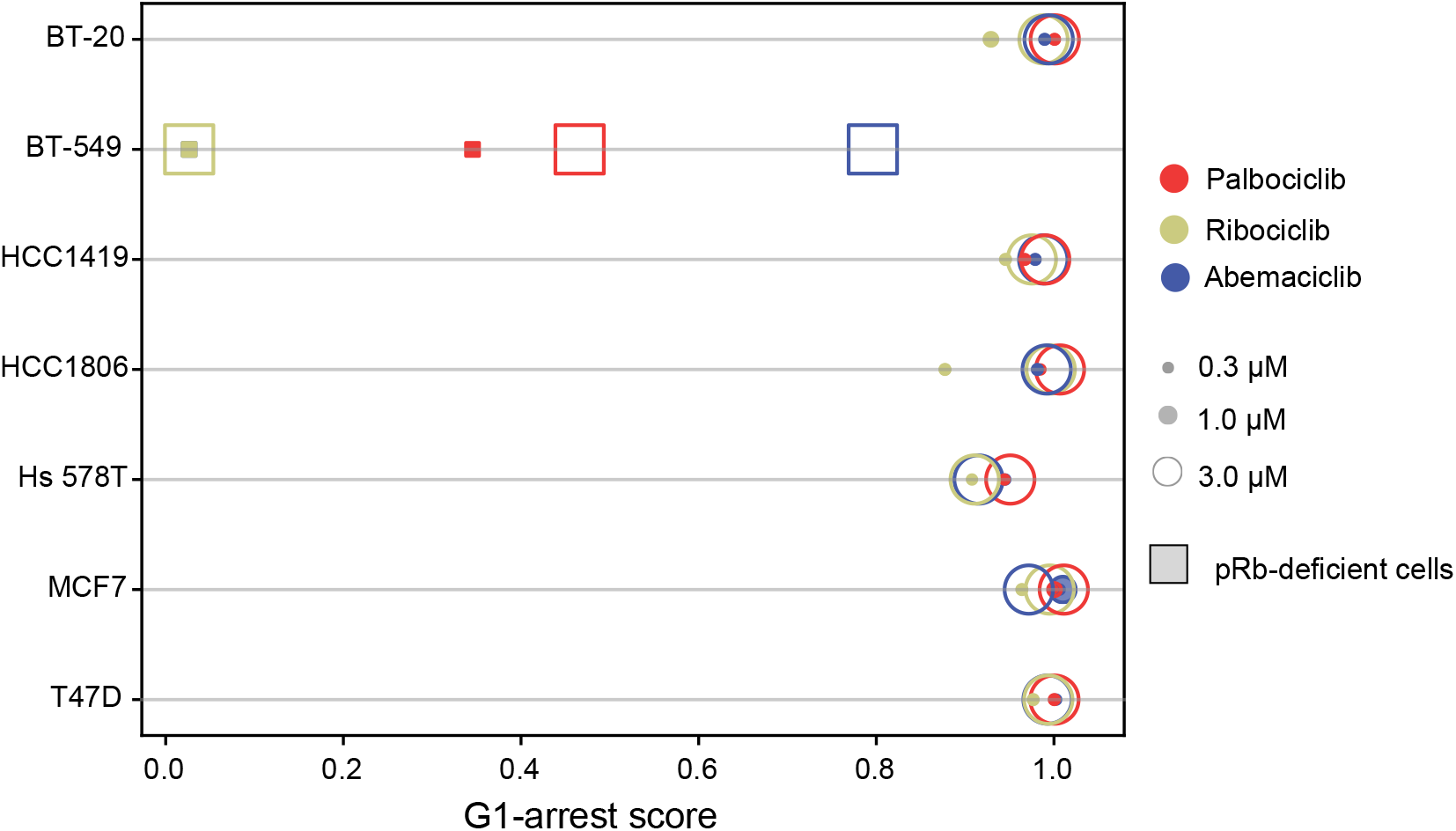
Related to Figure 1: G1-arrest transcriptional signature score. Score of the G1-arrest transcriptional signature per cell line following 6 hours of exposure to drug based on the RNA-seq data from Figure 1a.

**Figure S2.**
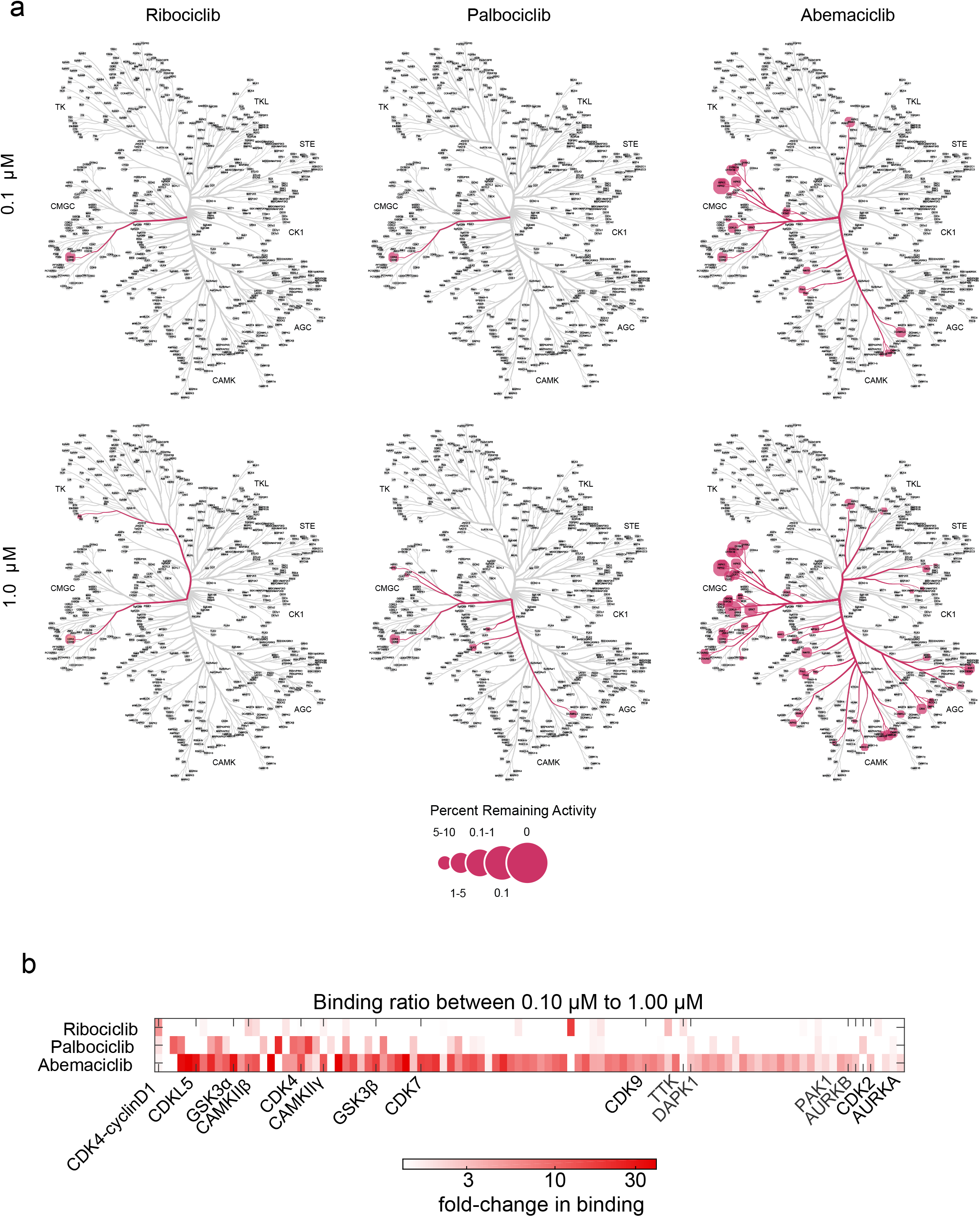
Related to Figure 3: KINOMEscan results for the three CDK4/6 inhibitors. **(a)** Kinases with less than 10% activity remaining for drug concentrations of 0.1 μM and 1.0 μM of each of the three CDK4/6 inhibitors (see Table S6). Images generated using Coral (Metz et al., 2018). **(b)** The differential binding between 0.1 μM and 1.0 μM of the 100 most bound kinases, plus kinases inferred to be inhibited by phosphoproteomics, for all drugs. CAMKIIα, CDK6, PKCγ, and PKCζ were not present in the panel.

**Figure S3.**
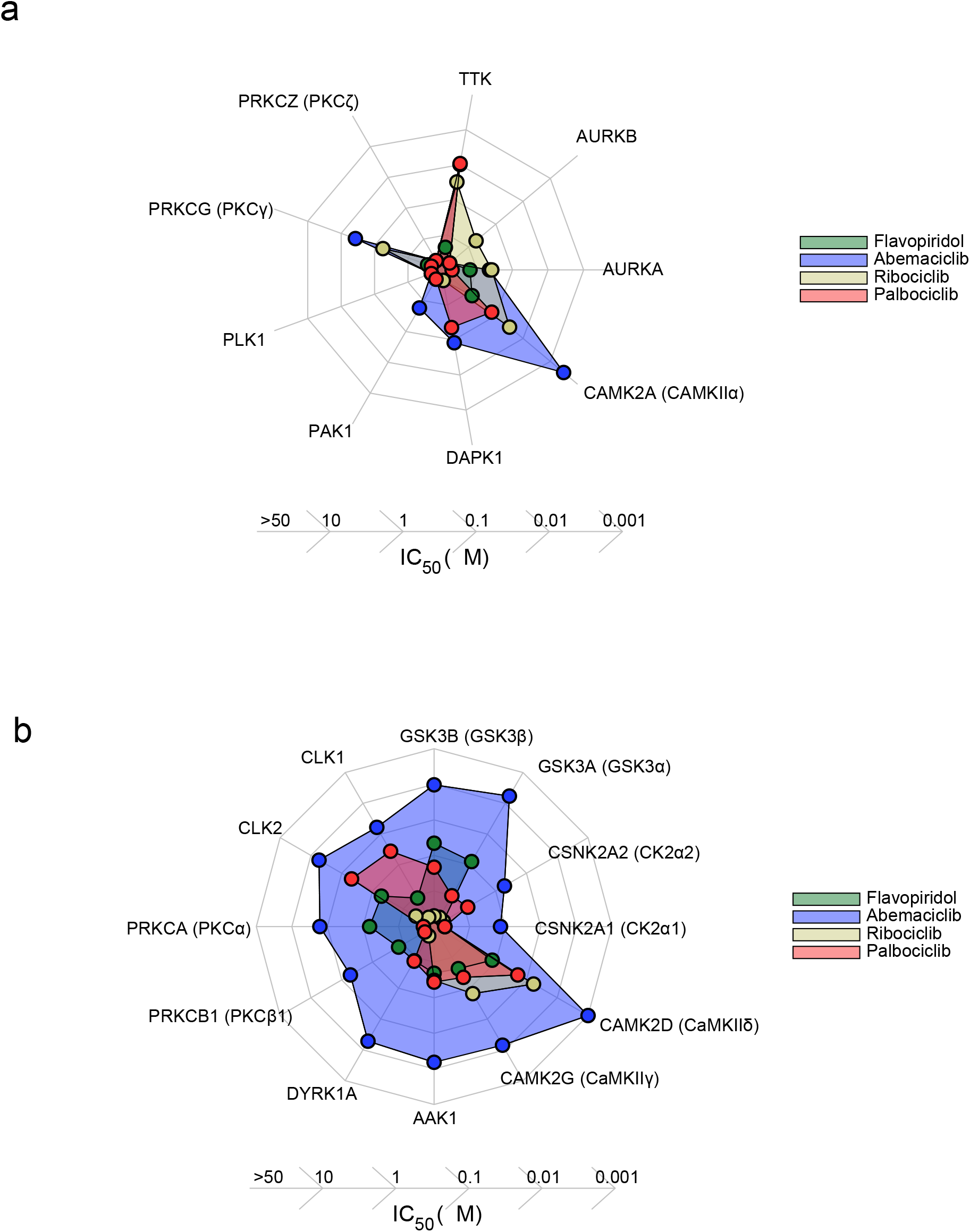
Related to Figure 3: *In vitro* activity of CDK4/6 inhibitors and alvocidib. **(a)** *IC_50_* values for CDK4/6 inhibitors and alvocidib for kinases other than CDKs inferred to be inhibited from phosphoproteomics data by either palbociclib or abemaciclib (see Figure 3b). **(b)** *IC_50_* values for CDK4/6 inhibitors and alvocidib for kinases inhibited by 10 μM abemaciclib in the MIB assay (log_2_ fold change > 2, only kinases non-overlapping with those shown in Figures 3e and S3a are shown).

**Figure S4.**
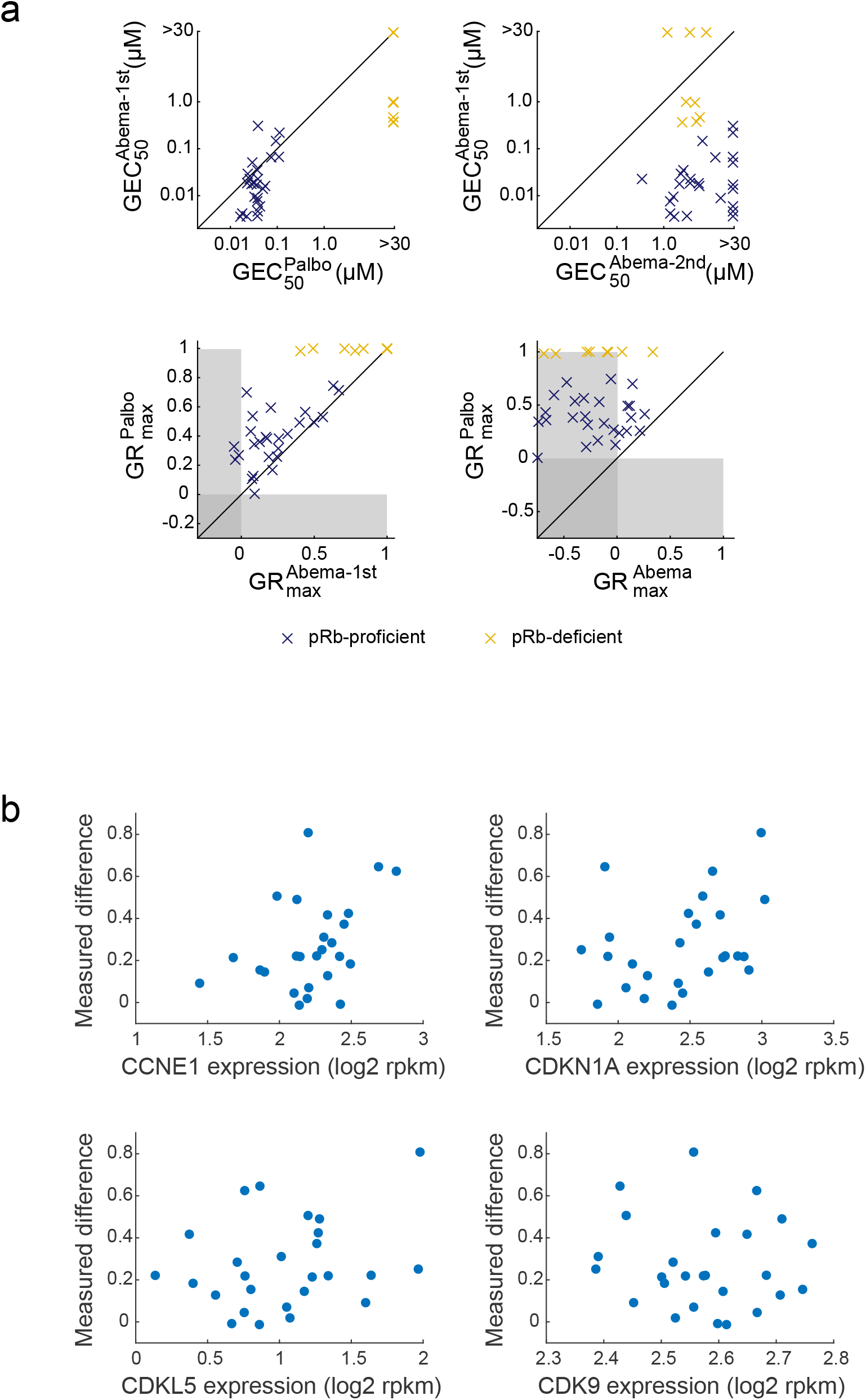
Related to Figure 4: Difference in efficacy between palbociclib and abemaciclib. **(a)** Comparison of fit parameters defined in Figure 4b for the dose response curves for palbociclib and abemaciclib shown in Figure 4a: (top) Mid-point concentrations for palbociclib (left) and the 2^nd^ phase of abemaciclib (right) versus the mid-point concentrations for the 1^st^ phase of abemaciclib. (bottom) Maximal efficacy for 1^st^ phase of abemaciclib (left) and abemaciclib (right) versus the maximal efficacy of palbociclib. **(b)** Measured difference in GR value between palbociclib and abemaciclib at 3 µM plotted against the variables of the predictive model defined in Figure 4c-d.

**Figure S5.**
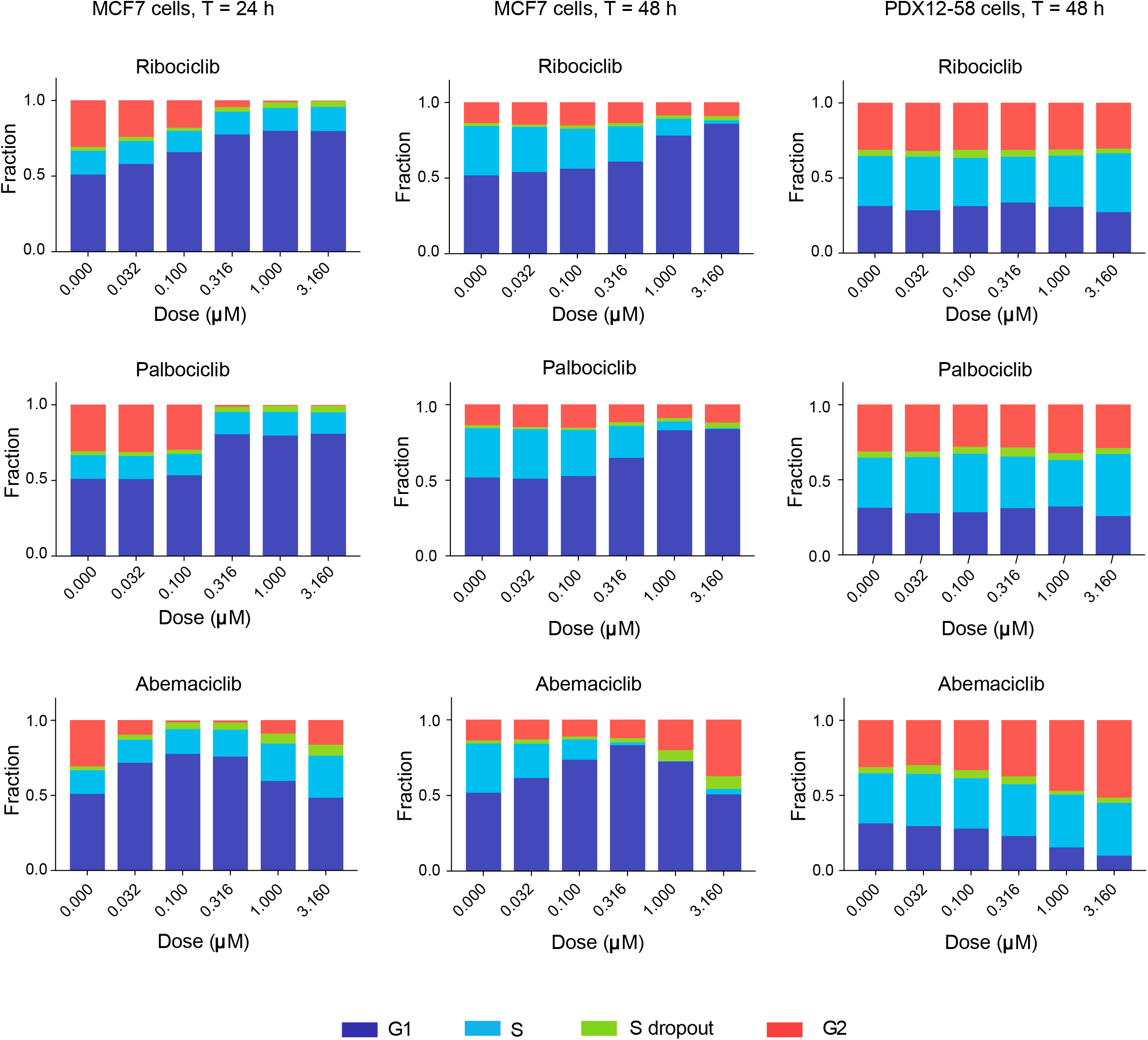
Related to Figure 5: Effect of CDK4/6 inhibitors on the distribution of cells through the cell cycle. Cell cycle distribution of MCF7 and PDX1258 breast cancer cell lines under the same conditions as shown in Figure 5. MCF7 cells exposed to one of three CDK4/6 inhibitors over a range of concentrations for 24 (left) or 48 (middle) hours, and in PDX-1258 cells, which are pRb-deficient in the same conditions for 48 hours (right). Cells were labeled with EdU to quantitate those in S-phase.

**Figure S6.**
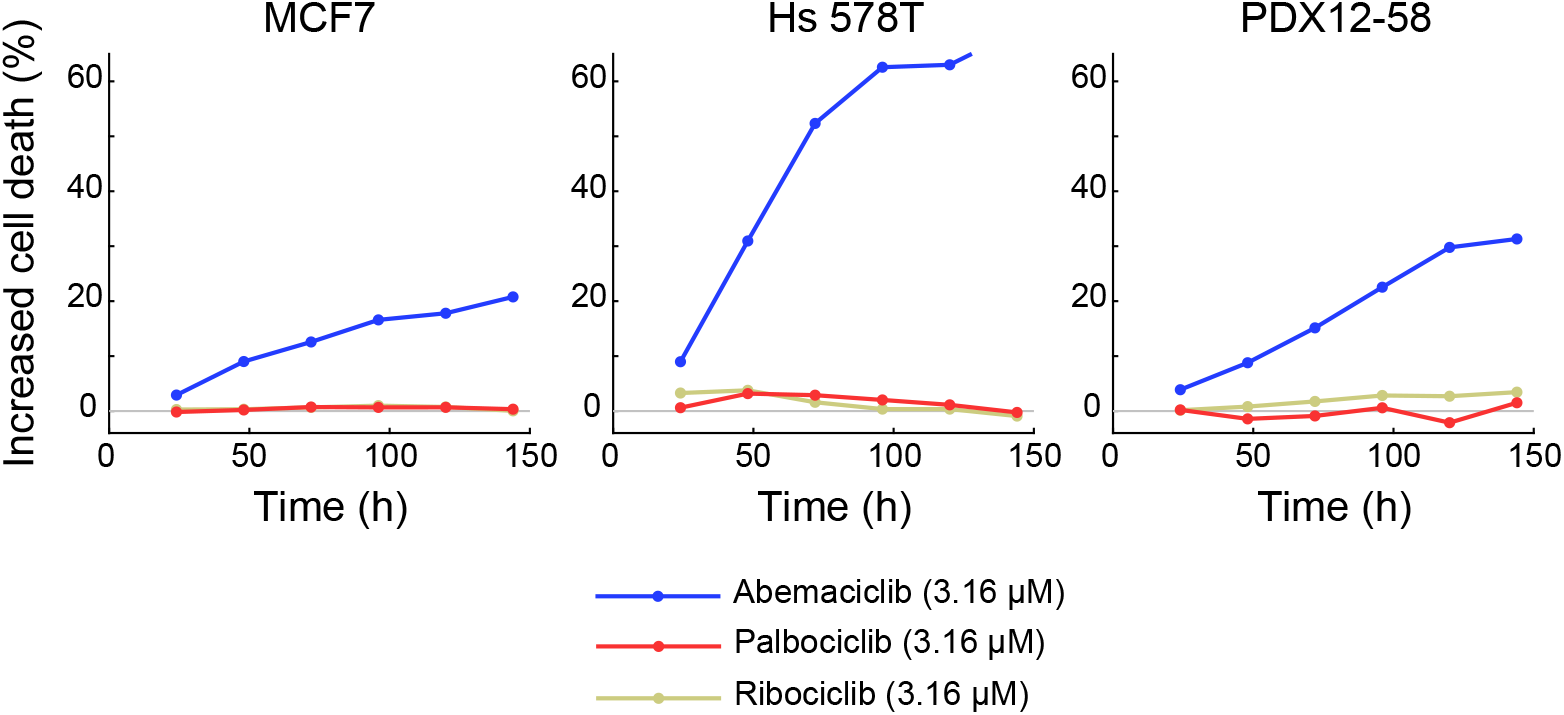
Related to Figure 7: Time dependent increase in the fraction of dead cells treated with CDK4/6 inhibitors. Increased percent of dead cells treated with 3.16 μM ribociclib, palbociclib, or abemaciclib relative to control conditions over time (see Figure 7a).

**Figure S7.**
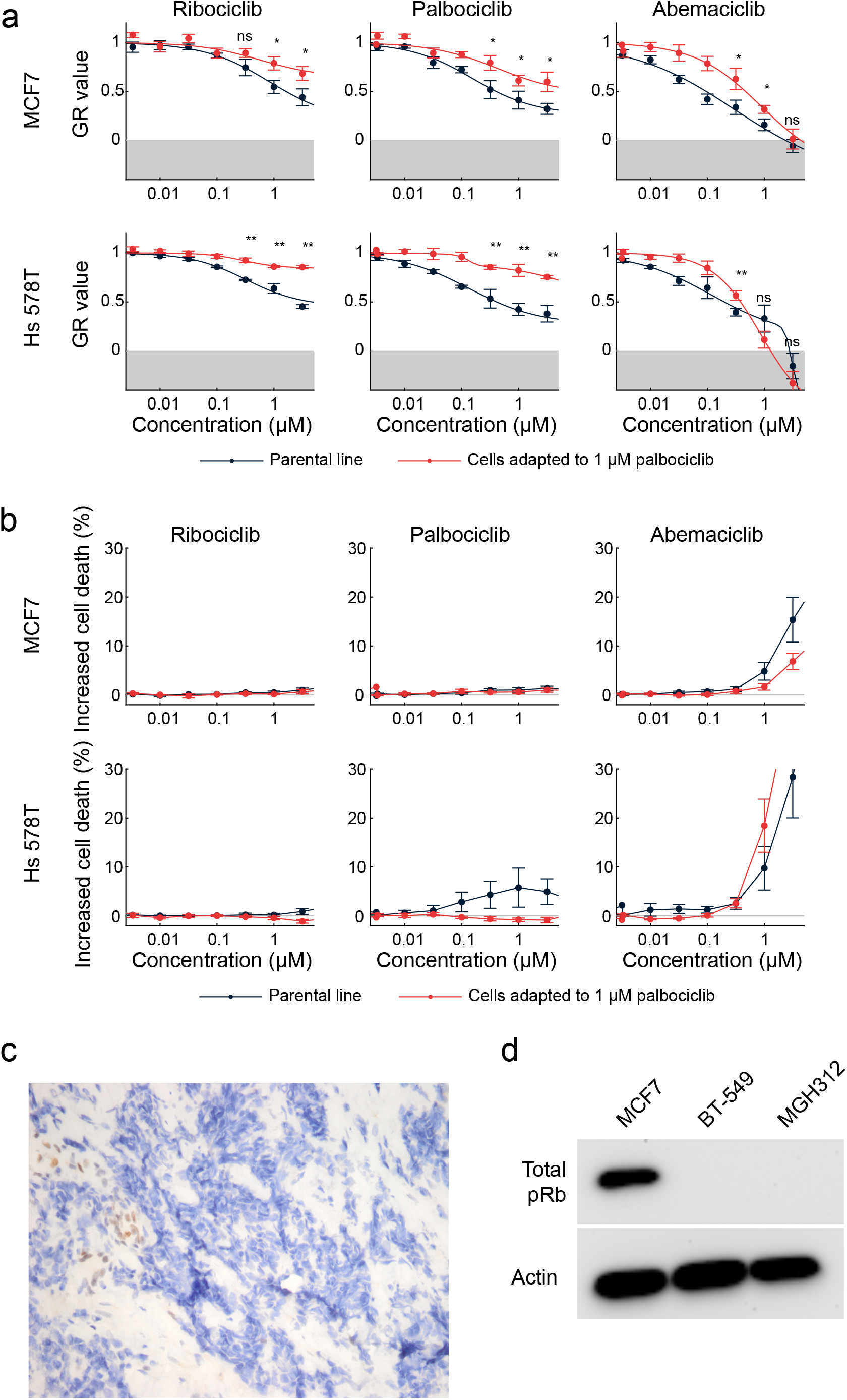
Related to Figure 7: Characterization of CDK4/6-inhibitor resistant cell lines. **(a)** GR values for cell lines adapted to grow in 1 μM palbociclib and their parental lines in response to 72-hour treatments with ribociclib, palbociclib, or abemaciclib. Error bars show the SEM of six replicates. **(b)** Increased percent of dead cells over vehicle-only control conditions for cell lines adapted to grow in 1 μM palbociclib and their parental lines in response to 72-hour treatments with ribociclib, palbociclib, or abemaciclib. Error bars show the SEM of six replicates. **(c)** pRb immunohistochemistry staining of the patient biopsy at the site from which the cell line MGH312 was derived. **(d)** Western Blot of total pRb for MCF7, BT-549, and MGH312 cells.

