## Supplementary material for "Multi-omics profiling establishes the polypharmacology of FDA Approved CDK4/6 inhibitors and the potential for differential clinical activity": Materials and Methods

### CONTACT FOR REAGENT AND RESOURCE SHARING

### EXPERIMENTAL MODEL AND SUBJECT DETAILS

#### Cell lines

All cell lines used in this study were of female human breast cancer origin except the MCF 10A and HME1 cell lines that were derived from non-transformed human breast epithelia. Cell lines were maintained, free of mycoplasma, in their recommended growth conditions listed below, and were identity-validated by STR profiling (Reid *et al*., 2004).

| *Cell line* | *Growth media* | *Growth conditions* |
|---|---|---|
| **BT20** | EMEM + 10% FBS + 1% P/S | 37˚C, 5% CO2 |
| **BT549** | RPMI-1640 + 10% FBS + 1% P/S, 1 ug/ml IN | 37˚C, 5% CO2 |
| **CAL120** | DMEM + 10% FBS + 1% P/S | 37˚C, 5% CO2 |
| **CAL51** | DMEM + 20% FBS + 1% P/S | 37˚C, 5% CO2 |
| **CAL851** | DMEM + 10% FBS + 1% P/S | 37˚C, 5% CO2 |
| **CAMA1** | EMEM + 10% FBS + 1% P/S | 37˚C, 5% CO2 |
| **HCC1143** | RPMI-1640 + 10% FBS + 1% P/S | 37˚C, 5% CO2 |
| **HCC1395** | RPMI-1640 + 10% FBS + 1% P/S | 37˚C, 5% CO2 |
| **HCC1419** | RPMI-1640 + 10% FBS + 1% P/S | 37˚C, 5% CO2 |
| **HCC1428** | RPMI-1640 + 10% FBS + 1% P/S | 37˚C, 5% CO2 |
| **HCC1500** | RPMI-1640 + 10% FBS + 1% P/S | 37˚C, 5% CO2 |
| **HCC1806** | RPMI-1640 + 10% FBS + 1% P/S | 37˚C, 5% CO2 |
| **HCC1937** | RPMI-1640 + 10% FBS + 1% P/S | 37˚C, 5% CO2 |
| **HCC1954** | RPMI-1640 + 10% FBS + 1% P/S | 37˚C, 5% CO2 |

| | | |
|---|---|---|
| **HCC38** | RPMI-1640 + 10% FBS + 1% P/S | 37˚C, 5% CO2 |
| **HCC70** | RPMI-1640 + 10% FBS + 1% P/S | 37˚C, 5% CO2 |
| **HME1** | MEMB + Lonza CC-3150 kit | 37˚C, 5% CO2 |
| **HS578T** | DMEM + 10% FBS + 1% P/S | 37˚C, 5% CO2 |
| **MCF10A** | DMEM/F12 (1:1) + 5% HS + 1% P/S, 20ng/ml EGF, 0.5mg/ml HC, 10 ug/ml IN, 100ng/ml CT | 37˚C, 5% CO2 |
| **MCF7** | DMEM + 10% FBS + 1% P/S | 37˚C, 5% CO2 |
| **MDAMB157** | L-15 + 10% FBS + 1% P/S | 37˚C, no CO2 |
| **MDAMB231** | DMEM + 10% FBS + 1% P/S | 37˚C, 5% CO2 |
| **MDAMB361** | L-15 + 20% FBS + 1% P/S | 37˚C, no CO2 |
| **MDAMB436** | L-15 + 10% FBS + 1% P/S, 10ug/ml IN | 37˚C, no CO2 |
| **MDAMB453** | L-15 + 10% FBS + 1% P/S | 37˚C, no CO2 |
| **MDAMB468** | L-15 + 10% FBS + 1% P/S | 37˚C, no CO2 |
| **MGH312** | RPMI-1640 + 10% FBS + 1% P/S | 37˚C, 5% CO2 |
| **PDX1258** | DMEM/F12 (3:1) + 7.5% FBS + 1% P/S, 0.125ng/ml EGF, 25ng/ml HC, 5ug/ml IN, 8.6ng/ml CT, 5 uM Y-27632 (Palechor-Ceron *et al*., 2013) | 37˚C, 5% CO2 |
| **PDX1328** | DMEM/F12 (3:1) + 7.5% FBS + 1% P/S, 0.125ng/ml EGF, 25ng/ml HC, 5ug/ml IN, 8.6ng/ml CT, 5 uM Y-27632 (Palechor-Ceron *et al*., 2013) | 37˚C, 5% CO2 |
| **PDXHCI002** | DMEM/F12 (3:1) + 7.5% FBS + 1% P/S, 0.125ng/ml EGF, 25ng/ml HC, 5ug/ml IN, 8.6ng/ml CT, 5 uM Y-27632 (Palechor-Ceron *et al*., 2013) | 37˚C, 5% CO2 |
| **SKBR3** | McCoy's + 10% FBS + 1% P/S | 37˚C, 5% CO2 |
| **SUM1315** | F-12 + 5% FBS + 1% P/S, 10ng/ml EGF, 5ug/ml IN, 10mM HEPES | 37˚C, 5% CO2 |
| **SUM149** | F-12 + 5% FBS + 1% P/S, 1ug/ml HC, 5ug/ml IN, | 37˚C, 5% CO2 |

| | | |
|---|---|---|
| | 10mM HEPES | |
| **SUM159** | F-12 + 5% FBS + 1% P/S, 1ug/ml HC, 5ug/ml IN, 10mM HEPES | 37˚C, 5% CO2 |
| **T47D** | RPMI-1640 + 10% FBS + 1% P/S, 1 ug/ml IN | 37˚C, 5% CO2 |

Abbreviations: fetal bovine serum (FBS), penicillin/streptomycin (P/S), insulin (IN), hydrocortisone (HC), epidermal growth factor (EGF), cholera toxin (CT). Reagent details can be found in the Key Resources Table.

##### Animals

Seven-week-old female NU/NU nude (Crl:NU-Foxn1$^{nu}$) mice (RRID IMSR_CRL:088) were used for this study (Charles River, Wilmington, MA). The animals were housed five per cage in the Harvard Center for Comparative Medicine animal facility and had *ad libitum* access to food and water (supplemented with 8 μg/ml 17 ß-estradiol to sustain growth of the hormone receptor positive xenografted tumor cells). Once tumors reached 250 mm$^3$ the mice were randomly assigned to treatment groups. All animal experiments were conducted in accordance with protocols approved by the Institutional Animal Care and Use Committee (IACUC) at Harvard Medical School.

#### METHOD DETAILS

**Dose response measurements**

Cells were plated at densities ranging from 500 to 2000 cells per well in 384-well Cell Carrier plates (Perkin Elmer, Waltham, MA) using a Multidrop Combi Reagent Dispenser (Thermo Fisher Scientific, Waltham, MA) and grown for 36 hours. Cells were treated with a dilution series of the indicated drugs by pin transfer or using a D300 Digital Dispenser (Hewlett-Packard, Palo Alto, CA). Drugs were obtained from commercial vendors and tested for purity in-house as described in detail in the HMS LINCS drug collection database (http://lincs.hms.harvard.edu/db/sm/). Cells were stained and fixed for analysis at the time of drug delivery and after 24 to 144 hours of incubation depending on the experiment. Cells were stained at the indicated time points with 2 μg/ml

Hoechst 33342 (Sigma Aldrich, St. Louis, MO) and 1:1000 LIVE/DEAD Far Red Dead Cell Stain (Thermo Fisher Scientific, Waltham, MA) for 30 minutes and fixed with 3.7% formaldehyde (Sigma Aldrich, St. Louis, MO) for 30 minutes. Fixed cells were imaged with a 10x objective using an Operetta microscope and analyzed using the Columbus image data storage and analysis system (Perkin Elmer, Waltham, MA). For most experiments, each condition was tested across three replicate plates and at least four wells per cell line per plate were untreated.

Nuclei counts were normalized to DMSO-treated controls on the same plate to yield relative cell count and normalized growth rate inhibition (GR) values for each technical replicate for each condition (Hafner *et al.*, 2016). Technical replicates were averaged to yield mean relative cell counts and the mean GR value for each condition within each biological replicate. Within each biological replicate, mean GR values for a given cell line / small molecule combination across all tested concentrations were fitted to a biphasic sigmoidal curve with the equation:

$$GR(c) = 2^{\log_2\left(GR_{max}^{1st} + \frac{1-GR_{max}^{1st}}{1+\left(c/GEC_{50}^{1st}\right)^{h^{1st}}} + 1\right) \cdot \log_2\left(GR_{max}^{2nd} + \frac{1-GR_{max}^{2nd}}{1+\left(c/GEC_{50}^{2nd}\right)^{h^{2nd}}} + 1\right)} - 1,$$

or with a single sigmoidal curve with the equation:

$$GR(c) = GR_{max} + \frac{1 - GR_{max}}{1 + \left(c/GEC_{50}\right)^{h}} + 1,$$

or with a flat line with the equation $GR(c) \equiv GR_{max}$. The significance of each curve was assessed using an F-test and the most complex model with $P < 0.05$ was considered to best fit the data. The parameters of the sigmoidal curve and the first phase of the biphasic curve are constrained as described in Hafner *et al.* (Hafner *et al.*, 2017). In the biphasic curve, the parameter $GEC_{50}^{2nd}$ is constrained to be above 0.3 $\mu$M. The time-dependent GR values (Hafner *et al.*, 2016) for Figure 7a were evaluated over a 48-hour interval.

**Phospho-pRb immunofluorescence and cell cycle analysis**

Cells were seeded in 384-well plates, allowed to adhere for 24-36 hours, treated with CDK4/6 inhibitors, incubated for the desired amount of time then fixed in 4%

formaldehyde, permeabilized with 0.5% Triton X-100 in PBS, and blocked with Odyssey blocking buffer (LI-COR, Lincoln, NE). Cells were labeled overnight at 4°C with a 1:800 dilution of anti-phospho-pRb Alexa-555 (Ser807/811) (Cell Signaling Technologies, Danvers, MA) and 2 μg/ml Hoechst 33342 (Sigma Aldrich, St. Louis, MO) prepared in Odyssey blocking buffer. Images were acquired with a Perkin Elmer Operetta microscope as described for the dose response measurements. Nuclei were segmented using Columbus software (Perkin Elmer, Waltham, MA) based on their Hoechst signal. DNA content was defined by the total Hoechst intensity within the nuclear mask. The average phospho-pRb intensity within the nuclear mask was determined, and a threshold for positivity was set by visually inspecting images of several control and treated wells per cell line. For more detailed cell cycle analysis, cells were labeled with a Click-iT™ EdU Alexa Fluor™ 488 Imaging Kit according to the manufacturer's directions (one hour EdU pulse prior to fixation) (Thermo Fisher Scientific, Waltham, MA). Imaging and segmentation were the same as for the immunofluorescence experiments. The average EdU intensity within the nuclear mask was determined. DNA content was used to identify cells in the G1 and G2 phases of the cell cycle, EdU intensity was used to identify cell in S phase. Cells with no EdU signal, and intermediate DNA content were classified as S-phase dropout cells.

**mRNA-seq**

Cells were seeded in 12-well plates, and allowed to adhere for 24 hours at which time CDK4/6 inhibitors were added. Cells were lysed in the plates after 6 or 24 hours, and RNA was extracted using Applied Biosystems MagMax 96 total RNA isolation kit (Thermo Fisher Scientific, Waltham, MA) with DNAse digestion according to the manufacturer's protocol. RNA was checked for quantity with a NanoDrop (Thermo Fisher Scientific, Waltham, MA) and for quality using an Agilent Bioanalyzer instrument (with RIN value > 9.0). Libraries were prepared using a TruSeq Stranded mRNA sample preparation kit (Illumina, San Diego, CA) from 500 ng of purified total RNA according to the manufacturer's protocol in a reduced reaction volume. The finished cDNA libraries were assessed for quality using a Bioanalyzer and quantified with a Quant-iT dsDNA Assay kit (Thermo Fisher Scientific, Waltham, MA). The uniquely indexed libraries were multiplexed based on this quantitation and the pooled sample was quantified by qPCR using the Kapa Biosystems (Wilmington, MA) library quantification kit by the Molecular Biology Core Genomics Facility at the Dana-Farber Cancer Institute and sequenced on a single Illumina NextSeq500 run with single-end 75bp reads.

Reads were processed to counts using the bcbio-Nextgen toolkit version 1.0.3a (https://github.com/chapmanb/bcbio-nextgen) as follows: (1) Reads were trimmed and clipped for quality control in cutadapt v1.12; (2) Read quality was checked for each sample using FastQC 0.11.5; (3) High-quality reads were then aligned into BAM files through STAR 2.5.3a using the human assembly GRCh37; (4) BAM files were imported into DEXSeq-COUNT 1.14.2 and raw counts TPM and RPKM were calculated. R package edgeR (Robinson, McCarthy and Smyth, 2010) 3.18.1 (R version 3.2.1) was used for differential analysis and generate log fold change, *P*-value and FDR.

**3’DGE sequencing**

Cells were plated at densities ranging from 500 to 2000 cells per well in a 384-well Cell Carrier plate (Perkin Elmer, Waltham, MA) and allowed to adhere for 24 hours. Cells were treated with the CDK4/6 inhibitors, alvocidib, or DMSO using a D300 Digital Dispenser (Hewlett-Packard, Palo Alto, CA). After six hours, the cells were washed once with PBS using an EL405x plate washer (BioTek, Winooski, VT), 10 $\mu$l of 1X TCL lysis buffer with 1% (v/v) ß-mercaptoethanol (Qiagen, Hilden, Germany) was added per well, and the plates were stored at -80˚C until the RNA extraction was performed. For RNA extraction, the cell lysate plate was thawed, vortexed briefly, and centrifuged for 1 min at 1000 rpm. Using a BRAVO (Agilent, Santa Clara, CA) liquid handler, the lysate was mixed thoroughly before transferring 10 $\mu$l to a 384 well PCR plate. 28 $\mu$l of SPRI beads (Beckman Coulter Genomics, Chaska, MN) were added directly to the lysate, mixed and incubated for 5 min. The plate was transferred to a magnetic rack to aggregate the beads, and incubated for 5 min prior to removing the liquid. The beads were washed with 80% ethanol twice, allowed to dry for 1 min, 20 $\mu$l of nuclease free water was added per well, the plate was removed from the magnetic rack and the beads were thoroughly resuspended. Following a 5 min incubation, the plate was returned to the magnetic rack and incubated an additional 5 min before transferring the supernatant to a fresh PCR plate. 5 $\mu$l of the supernatant was transferred to a separate plate containing RT master mix and 3' and 5' adapters for reverse transcription and template switching (Soumillon *et al.*, 2014), and incubated for 90 min at 42˚C. The cDNA was pooled and purified with a QIAquick PCR purification kit according to the manufacturer's directions with the final elution in 24 $\mu$l of nuclease free water. This was followed by an exonuclease I treatment for 30 min at 37˚C that was stopped with a 20 min incubation at 80˚C. The cDNA was then amplified using the Advantage 2 PCR Enzyme System (Takara, Fremont, CA) for 5 cycles, and purified using AMPure XP magnetic beads (Beckman Coulter Genomics, Chaska, MN). Library preparation was completed with 55 ng input using a Nextera DNA kit (Illumina, San Diego, CA) following the manufacturer's instructions, amplified 5 cycles, and purified with AMPure XP magnetic beads (Beckman Coulter Genomics, Chaska, MN). A Pippin PREP purification of the sample from 300-800bp was performed, it was then quantified by qPCR and sequenced on a single Illumina NextSeq run with 75bp paired end reads at the Harvard University Bauer Core Facility.

Reads were processed to counts through the bcbio-nextgen single cell/DGE RNA-seq analysis pipeline (https://bcbio-nextgen.readthedocs.io/en/latest/contents/pipelines.html) a brief description follows: The well barcode and UMIs were identified for all reads and all reads not within one edit distance of a known well barcode were discarded. Each surviving read was quasialigned to the transcriptome (GRCh38) using RapMap (Srivastava *et al.*, 2016). Reads per well were counted using UMIs (Svensson *et al.*, 2017), discarding duplicated UMIs, weighting multimapped reads by the number of transcripts they aligned to and collapsing counts to genes by adding all counts for each transcript of a gene. The R package edgeR 3.18.1 (R version 3.2.1) was used for differential expression analysis.

**Clustering analysis of the mRNA-seq data and L1000 signatures**

Differential gene expression signatures were clustered along samples and genes based on the cosine distance for the $\log_2$(fold-change) using MATLAB default functions. $\log_2$(fold-change) values for genes with FDR values above 0.2 were set to zero. In Figure 1a, the two down-regulated gene clusters were defined manually based on the dendrogram of the genes. The 'LINCS_L1000_Chem_Pert_down' library obtained from Enrichr (Kuleshov *et al*., 2016) was used as the reference signature of genes downregulated upon drug perturbation (Table S2). Enrichment analysis was performed on the two down-regulated gene clusters (Figure 1a) against the reference library using the GSEA algorithm (gsea2-2.2.3.jar from the Broad Institute (Subramanian *et al*., 2005)). Enrichment scores for 31 well-annotated drugs that feature in the library were reported (Figure 1b-c) as $-\log_{10}(P$-value). G1-arrest and pan-CDK scores for each condition (Figure S1, Figure 1d) were computed as the mean $\log_2$(fold-change) across the genes in each of the two red (G1-arrest) and cyan (pan-CDK) down-regulated gene clusters identified in Fig 1a. G1-arrest and pan-CDK scores for 3' DGEseq (Figure 2) and MCF7-xenograft mRNAseq (Figure 6b) were computed on the same set of down-regulated genes.

**Phosphoproteomics mass spectrometry**

MCF7 cells were treated with 0.3 $\mu$M and 3 $\mu$M of palbociclib or abemacicblib, or DMSO control for 1 hour in duplicate. For each sample, 4.5 mg of protein was utilized to perform serine and threonine phosphoproteome analysis. The samples were digested using Trypsin (Promega, Madison, WI), acidified and desalted using C18 Sep-Pak (Waters, Milford, MA). Phosphopeptides were enriched using the Thermo Scientific High-Select Fe-NTA Phosphopeptide Enrichment Kit. The samples were labeled using a TMT 10plex Mass Tag Labeling kit (Thermo Fisher Scientific, Waltham, MA) and the reaction was quenched by adding hydroxylamine to a final concentration of 0.5% (v/v) (Kettenbach and Gerber, 2011; Paulo *et al*., 2015). The sample was then enriched for phosphotyrosine-containing peptides using the pY-1000 antibody (Cell Signaling Technologies, Danvers, MA) coupled to Pierce Protein A Agarose beads (Thermo Fisher Scientific, Waltham, MA). The flow-through from the pY sample was kept and desalted for pS and pT analysis. 24 fractions (phosphoproteomics) were then desalted using the C18 StageTip procedure (Rappsilber, Mann and Ishihama, 2007). All MS analyses were performed on an Orbitrap Fusion Lumos mass spectrometer (Thermo Fisher Scientific, Waltham, MA) using a multi-notch MS3 method (Ting *et al*., 2011; McAlister *et al*., 2014). Raw data were converted to mzXML and searched via Sequest (Eng, McCormack and Yates, 1994) version 28 against a concatenated Uniprot database (downloaded 02/04/2014). Linear discriminate analysis was used to distinguish forward and reverse hits and reverse hits were filtered to an FDR of 1% at the protein level. Site localization confidence was assessed using the A-score method (Beausoleil *et al*., 2006). Reporter ion intensities were quantified and normalized as described earlier (Paulo *et al*., 2015).

**Annotation of phosphopeptides with upstream kinases**

16,300 phosphopeptides were detected across all conditions in MCF7 cells. The PhosphoSitePlus (PSP) database (Hornbeck *et al*., 2012), which contains curated annotations of upstream kinases, was queried using phosphopeptide sequence motifs and UniProt IDs as identifiers. Only ~6.3% of the phosphopeptides detected by phosphoproteomics had experimentally verifiable kinase annotations on PSP. The NetworKIN algorithm (Horn *et al*., 2014) that predicts upstream kinases, based on phosphopeptide sequences and STRING evidence, was used to identify kinases for the remaining phosphosites. A further 14% of phosphosites were annotated with predicted kinases (NetworKIN Score > 4). In total, 3145 phosphopeptides from 1242 proteins were annotated as being phosphorylated by 365 kinases (8297 kinase-peptide interaction pairs).

**Differential kinase activity score using GSEA**

Based on the method described previously (Drake *et al*., 2012), a custom python package was developed to infer differential kinase activity across drug treatments (https://github.com/datarail/msda). A kinase set library was assembled using the identified kinase-substrate relationships. The kinase set library is composed of kinases and their corresponding sets of phosphopeptide substrates. Only kinase sets that had more than 25 downstream phosphosites were used. The final kinase set library was composed of 60 kinases that phosphorylate 2597 peptides. For each phosphopeptide, the mean difference between the replicates and the maximum difference across conditions were computed. If the delta between the two scores was less than 1, then the phosphopeptide measurement was considered noisy and discarded, resulting in a final list of 9958 phosphopeptides (Table S4). For each of the four treatment conditions, the average log2 (fold-change) was computed relative to the untreated control. Using the phosphopeptide log2(fold-change) values as input and the final kinase set library, GSEA algorithm (gsea2-2.2.3.jar from Broad Institute (Subramanian *et al*., 2005)) was used to infer the enrichment score ($P < 0.05$ and FDR < 0.2). The enrichment score is a proxy metric for the differential activity of the kinases.

**Measurement of kinase inhibition with kinobeads**

Multiplex inhibitor beads (MIB) (Duncan *et al*., 2012) were generously provided by Gary Johnson (University of North Carolina). A mixed cell lysate comprised of K562, COLO0205, SK-N-BE(2), MV-4-11 cells was prepared as previously described (Médard *et al*., 2015), and clarified by filtration through 0.45 μm and 0.22 μm filters. 3 mg of the mixed cell lysate was treated with CDK4/6 inhibitors or DMSO overnight at 4˚C with continuous rocking. The samples were enriched for kinases by passing them through a sepharose bead column followed by a MIB column. The samples were washed with MIB wash buffer (50 mM HEPES pH 7.5, 0.5% Triton X-100, 1 mM EDTA, 1 mM EGTA) with high (1 M NaCl) and low (150 mM NaCl) salt, and then in low salt MIB buffer containing 0.1% SDS (w/v). Kinases bound to the MIBs were eluted twice with 500 μl/column of MIB elution buffer (0.5% (w/v) SDS, 10 μM DTT, 0.1 M Tris-HCl pH 6.8). The eluent was boiled for 15 min at 97˚C, and alkylated with 0.5 M iodoactamide (30 μl per ml of sample) for 30 min at room temperature. The samples were precipitated with tricholoracetic acid (25% final volume), washed twice with methanol, and dried. The samples were solubilized in 8 M urea in 20 mM EPPS. Additional EPPS was added to decrease the concentration of urea to 2 M prior to adding acetonitrile (ACN) and lysC (2 μg/μl) for 3 hours at room temperature. The samples were digested with trypsin (0.5 μg/μl) overnight at 37˚C. Additional ACN was added, followed by 5 μl of tandem mass tag (TMT) labels (Thermo Fisher Scientific, Waltham, MA) for 1 hour at room temperature. At this stage a ratio check was performed to ensure equal loading of each individually TMT-labeled sample, and to check the efficiency of the labeling reaction. The labeling reactions were quenched with 5 μl of 10% hydroxylamine for 10 min at room temperature, at which point the samples were pooled, diluted with 100% formic acid, evaporated to 0.5 ml, diluted with 1% formic acid, and then desalted by passing through a solid phase extraction cartridge (Thermo Fisher Scientific, Waltham, MA). Samples were eluted in 70% ACN and 1% formic acid, evaporated, and reconstituted in 300 μl of 0.1% TFA. The samples were analyzed on an Orbitrap Fusion Lumos mass spectrometer. Peptide intensities of the proteins pulled down by the MIBs were summed to obtain total protein intensities. The protein intensities were normalized using the iBAQ method (Schwanhäusser *et al*., 2011). For each treatment condition, log2(fold-change) values were computed relative to untreated (DMSO) control.

***In vitro* measurement of kinase inhibitory activity**

Ribociclib, palbociclib, and abemaciclib were assayed using the KINOMEscan® assay platform (DiscoverX, Fremont, CA). Data are reported as percent of remaining activity at either 0.1 or 1.0 $\mu$M drug concentration. The activity of ribociclib, palbociclib, abemaciclib, and alvocidib on multiple CDK-cyclin complexes and other kinases were assayed using Thermo Fisher Scientific SelectScreen Kinase Profiling service. The 'Adapta™' assay was used for CDK4/cyclin D1, CDK4/cyclin D3, CDK6/cyclin D1, CDK7/cyclin H/MNAT1, and CDK9/cyclin T1. The 'LanthaScreen™' Kinase Binding assay was used for CDK2/cyclin A1, CDK2/cyclin E1, CDK9/cyclin K, and TTK. The 'Z′-LYTE™' assay was used for CDK1/cyclin B, AURKA, AURKB, CAMK2A, GSK3B, and PLK1. The ATP concentration was Km[app] when available or 10 $\mu$M otherwise.

**Western blots**

20 $\mu$g of whole cell lysate (Figure 7b) or 12 $\mu$g of whole cell lysate (Figure S7), prepared in M-PER lysis buffer (Thermo Fisher Scientific, Waltham, MA) with complete protease inhibitor cocktail (Sigma Aldrich, St. Louis, MO), was added per well in Mini-PROTEAN TGX precast gels (Bio-Rad, Hercules, CA). Primary mouse monoclonal pRb, cyclin E, and β-actin antibodies were used at 1:1000 dilutions. Secondary anti-mouse IgG, HRP-linked was used 1:2000. All antibodies were from Cell Signaling Technologies (Danvers, MA).

**Immunohistochemistry**

A 4 $\mu$m slice of a formalin-fixed, paraffin-embedded, biopsy of the liver lesion from which the MGH312 cell line was derived was mounted on a standard glass slide and stained for RB expression using a Leica Bond autostainer. The primary Rb antibody (clone 1F8; Bio SB, Santa Barbara, CA) was diluted 1:500 in Leica Bond Diluent and incubated for 15 min. The slide was counterstained with hematoxylin.

**Identifying genes associated with differential efficacy of abemaciclib and palbociclib**

~~Using the baseline mRNA expression of~~ We selected 30 genes ~~linked~~ related to the cell cycle (cyclins, CDKs, CDKLs, and CDKNs), whether or not they are known CDK4/6 on- and off-targets. Using the baseline mRNA expression of these genes, we built a multilinear model (MATLAB function 'fitglm') to predict the difference in GR values at 3.16 $\mu$M between palbociclib and abemaciclib for the pRb-proficient cell lines profiled in Figure 4a. Predictors with non-significant coefficients ($P > 0.05$) were iteratively removed until only significant coefficients remained. A leave-one-out cross validation was performed with the remaining predictors to yield the results in Figure 4c. Note that results were qualitatively similar if the pRb-deficient cell lines were included.

**In vivo studies**

Thirty-five seven-week-old NU/NU nude mice (Charles River Laboratories, Wilmington, MA) were supplemented with 8 $\mu$g/ml 17ß-estradiol (Sigma Aldrich, St. Louis, MO) by adding it to their drinking water five days prior to tumor engraftment, and replacing it twice per week. Mice were engrafted with 5 x $10^6$ MCF-7 cells 1:1 in growth factor reduced matrigel (Corning, Corning, NY) subcutaneously in each flank, and allowed to grow to ~300 $mm^3$. The animals were then randomly assigned to treatment groups, and treated daily for four days with ribociclib (150 mg/kg), palbociclib (150 mg/kg), abemaciclib (25, 75, 100, 125, or 150 mg/kg), or vehicle control (0.5% (w/v) hydroxyethyl cellulose (Sigma Aldrich, St. Louis, MO) and 0.05% (v/v) antifoam (Sigma Aldrich, St. Louis, MO) in water) by oral gavage. Animals were sacrificed two hours after receiving the last dose. The tumors were excised and cut in half, one half was fixed in 4% formaldehyde at 4˚C and transferred to 0.1% sodium azide after 48 hours, the other flash frozen, and a thin slice from the center of the tumor was placed in RNA*later* at 4˚C (Qiagen, Hilden, Germany), after 48 hours the RNA*later* was aspirated, and samples were transferred to -80˚C.

The fixed tumor samples were paraffin-embedded at the Harvard Medical Area Rodent Histopathology Core and a tissue microarray (TMA) was constructed at the Tissue

Microarray & Imaging Core by arraying three 1 mm cores per sample in a block. Sequential 5 μm slices were mounted on superfrost slides. The slides were subjected to manual dewaxing and antigen retrieval as described previously (Lin *et al*., 2017). The slides were then blocked with Odyssey buffer (LI-COR, Lincoln, NE) and pre-stained with secondary antibodies prior to beginning cyclic immunofluorescence (Lin *et al*., 2017). The antibodies used in this study are listed in the Key Resources Table. Images were acquired on a RareCyte CyteFinder (Seattle, WA) slide scanning microscope with a 10X 0.3 NA objective. Image quantitation was performed in ImageJ as previously described (Lin *et al*., 2017). Human cells were distinguished from mouse cells based on e-cadherin and vimentin intensities, and only the e-cadherin-high, vimentin-low cells were included in subsequent analyses. A threshold for phospho-pRb positive cells was set manually by comparing the intensity distributions of phospho-pRb staining in tumors from mice that received the vehicle control and 150 mg/kg palbociclib.

The RNA*later* preserved samples were thawed on ice, 600 μl of RLT with 10% 2-mercaptoethanol was added and the tumors were manually dissociated with microfuge pestles (Thomas Scientific, Swedesboro, NJ). Samples were passed through QiaShredder columns, and then loaded on RNeasy columns (Qiagen, Hilden, Germany) and processed according to the manufacturer's specifications with a 30 min incubation in DNase. Library preparation, and analysis were performed as described in section 4. A single Illumina NextSeq500 run with single-end 75bp reads was performed at the Harvard Medical School Biopolymers Facility. Reads were processed as described in section 4, with the additional step that the alignment algorithm identified and excluded reads that aligned with the mouse genome to ensure that downstream analyses were performed on the xenograft transcripts only. Non-coding genes were excluded from the transcript per million (TPM) counts table and Principal Component Analyses (PCA) was performed. For each treated sample, the fold-change of transcripts relative to vehicle control was computed using edgeR (Robinson, McCarthy and Smyth, 2010). G1 and pan-CDK scores were computed as described in section 6.

#### QUANTIFICATION AND STATISTICAL ANALYSIS

Image quantification was performed with Columbus (Perkin Elmer, Waltham, MA) software. All subsequent analyses were performed using MATLAB and python. All relevant statistical details are included in the figure captions, and text. Additional details for each experiment type are included in the METHOD DETAILS section of the STAR Methods.

#### DATA AND SOFTWARE AVAILABILITY

The RNA sequencing data sets related to Figures 1, 2, and 6 have been deposited on GEO, and can be found under accession numbers GSE99116, GSE125215 and GSE124854 respectively. The phosphoproteomics data set related to Figure 3 is freely available on Synapse, ID syn11622501, https://www.synapse.org/#!Synapse:syn11622501. The dose response data sets related to Figure 4 are available in the HMS LINCS database, IDs 20343 and 20344, http://lincs.hms.harvard.edu/db/datasets/20343/ http://lincs.hms.harvard.edu/db/datasets/20344/.

#### KEY RESOURCES TABLE

| REAGENT or RESOURCE | SOURCE | IDENTIFIER |
|---|---|---|
| Antibodies | | |
| Phospho-pRb (Ser807/811) (clone D20B12) Alexa 555 | Cell Signaling Technologies | Cat # 8957; RRID AB_2728827 |
| pRb (clone 4H1) | Cell Signaling Technologies | Cat # 9309; RRID AB_823629 |
| Cyclin E1 (clone HE12) | Cell Signaling Technologies | Cat # 4129; RRID AB_2071200 |
| β-Actin (clone 8H10D10) | Cell Signaling Technologies | Cat # 3700; RRID AB_10985704 |
| Anti-mouse IgG, HRP-linked | Cell Signaling Technologies | Cat # 7076; RRID AB_330924 |
| Vimentin (clone D21H3) Alexa 555 | Cell Signaling Technologies | Cat # 9855; RRID AB_10859896 |
| E-cadherin (clone 24E10) Alexa 488 | Cell Signaling Technologies | Cat # 3199; RRID AB_823441 |
| Chemicals, Peptides, and Recombinant Proteins | | |

| | | |
|---|---|---|
| Palbociclib | MedChem Express | Cat # HY-50767, batch # 16349 |
| Abemaciclib | MedChem Express | Cat # HY-16297, batch # 08492 |
| Ribociclib | MedChem Express | Cat # HY-15777, batch # 11003 |
| Alvocidib | Haoyuan chemexpress | Cat # HY-10005, batch # HY-009_TM-20090429 |
| Fetal bovine serum | Life Technologies | 26140-079 |
| Horse Serum | Life Technologies | 16050-122 |
| Penicillin/Streptomycin | Corning | 30-002-Cl |
| Epidermal growth factor | PeproTech | AF-100-15 |
| Insulin | Sigma Aldrich | I1882 |
| Hydrocortisone | Sigma Aldrich | H0888 |
| Cholera toxin | Sigma Aldrich | C8052 |
| Y-27632 | Enzo Life Sciences | ALX-270-333-M025 |
| Critical Commercial Assays | | |
| KINOMEscan | DiscoverX | SCANmax |
| SelectScreen Z' lyte | Life Technologies | Z'Lyte |
| SelectScreen Lantha | Life Technologies | Lantha |
| SelectScreen Adapta | Life Technologies | Adapta |
| TruSeq kit | Illumina | Cat # 20019792 |
| Deposited Data | | |
| mRNAseq on cell lines | This paper | GEO GSE99116 |
| Phosphoproteomics | This paper | Synapse syn11622501 |
| Dose response | This paper | LINCS DB 20343, 20344 |
| 3' DGEseq on cell lines | This paper | GEO GSE125215 |
| mRNAseq on xenografts | This paper | GEO GSE124854 |
| Experimental Models: Cell Lines | | |
| BT20 | ATCC | HTB-19; RRID CVCL_0178 |
| BT549 | ATCC | HTB-122; RRID CVCL_1092 |
| CAL120 | DSMZ | ACC 459; RRID CVCL_1104 |
| CAL51 | DSMZ | ACC 302; RRID CVCL_1110 |
| CAL851 | DSMZ | ACC 440; RRID CVCL_1114 |

| | | |
|---|---|---|
| CAMA1 | ATCC | HTB-21; RRID CVCL_1115 |
| HCC1143 | ATCC | CRL-2321; RRID CVCL_1245 |
| HCC1395 | ATCC | CRL-2324; RRID CVCL_1249 |
| HCC1419 | ATCC | CRL-2326; RRID CVCL_1251 |
| HCC1428 | ATCC | CRL-2327; RRID CVCL_1252 |
| HCC1500 | ATCC | CRL-2329; RRID CVCL_1254 |
| HCC1806 | ATCC | CRL-2335; RRID CVCL_1258 |
| HCC1937 | ATCC | CRL-2336; RRID CVCL_0290 |
| HCC1954 | ATCC | CRL-2338; RRID CVCL_1259 |
| HCC38 | ATCC | CRL-2314; RRID CVCL_1267 |
| HCC70 | ATCC | CRL-2315; RRID CVCL_1270 |
| HME1 | ATCC | CRL-4010; RRID CVCL_3383 |
| HS578T | ATCC | HTB-126; RRID CVCL_0332 |
| MCF10A | ATCC | CRL-10317; RRID CVCL_0598 |
| MCF7 | ATCC | HTB-22; RRID CVCL_0031 |
| MDAMB157 | ATCC | HTB-24; RRID CVCL_0618 |
| MDAMB231 | ATCC | HTB-26; RRID CVCL_0062 |
| MDAMB361 | ATCC | HTB-27; RRID CVCL_0620 |
| MDAMB436 | ATCC | HTB-130; RRID CVCL_0623 |
| MDAMB453 | ATCC | HTB-131; RRID CVCL_0418 |
| MDAMB468 | ATCC | HTB-132; RRID CVCL_0419 |
| MGH312 | MGH (Crystal *et al*., 2014) | |
| PDX1258 | Brugge lab | |
| PDX1328 | Sorger lab | |

| | | |
|---|---|---|
| PDXHCI002 | Brugge lab | |
| SKBR3 | ATCC | HTB-30; RRID CVCL_0033 |
| SUM1315 | University of Michigan | SUM-1315MO2; RRID CVCL_5589 |
| SUM149 | Asterand | SUM-149PT; RRID CVCL_3422 |
| SUM159 | Asterand | SUM-159PT; RRID CVCL_5423 |
| T47D | ATCC | HTB-133; RRID CVCL_0553 |
| Software and Algorithms | | |
| MATLAB (R2016b) | MathWorks | https://www.mathworks.com/products/matlab.html |
| Columbus (v2.7.0) | Perkin Elmer, Waltham, MA | http://www.perkinelmer.com/product/image-data-storage-and-analysis-system-columbus |
| bcbio-Nextgen toolkit (v1.0.3a) | | https://github.com/chapmanb/bcbio-nextgen |
| edgeR v3.18.1 (R v3.2.1) | (Robinson, McCarthy and Smyth, 2010; McCarthy, Chen and Smyth, 2012) | https://bioconductor.org/packages/release/bioc/html/edgeR.html |
| Sequest (v28) | (Eng, McCormack and Yates, 1994) | http://fields.scripps.edu/yates/wp/ |
| Kinase activity inference | | https://github.com/datarail/msda |
